# Gasdermin-E mediates mitochondrial damage in axons and neurodegeneration

**DOI:** 10.1101/2022.11.01.513927

**Authors:** Dylan V Neel, Himanish Basu, Georgia Gunner, Matthew D Bergstresser, Richard M. Giadone, Haeji Chung, Rui Miao, Vicky Chou, Eliza M. Brody, Xin Jiang, Edward B. Lee, Christine Marques, Aaron Held, Brian Wainger, Clotilde Lagier-Tourenne, Yong-Jie Zhang, Leonard Petrucelli, Tracy L. Young-Pearse, Alice S Chen-Plotkin, Lee L. Rubin, Judy Lieberman, Isaac M Chiu

## Abstract

Mitochondrial dysfunction and axon loss are hallmarks of neurologic diseases. Gasdermin (GSDM) proteins are executioner pore-forming molecules that mediate cell death, yet their roles in the central nervous system (CNS) are not well understood. Here, we find that one GSDM family member, GSDME is expressed by both mouse and human neurons. GSDME plays a role in mitochondrial damage and axon loss. Mitochondrial neurotoxins induced caspase-dependent GSDME cleavage and rapid localization to mitochondria in axons, where GSDME promoted mitochondrial depolarization, trafficking defects, and neurite retraction. The frontotemporal dementia (FTD)/amyotrophic lateral sclerosis (ALS)-associated proteins TDP-43 and PR-50 induced GSDME-mediated damage to mitochondria and neurite loss. GSDME deficiency prolonged survival, ameliorated motor dysfunction, and rescued motor neuron loss in the SOD1^G93A^ mouse model of ALS. GSDME knockdown also protected against neurite loss in ALS patient iPSC-derived motor neurons. Thus, we identify GSDME as an executioner of neuronal mitochondrial dysfunction that contributes to neurodegeneration.

**Graphical Abstract:** 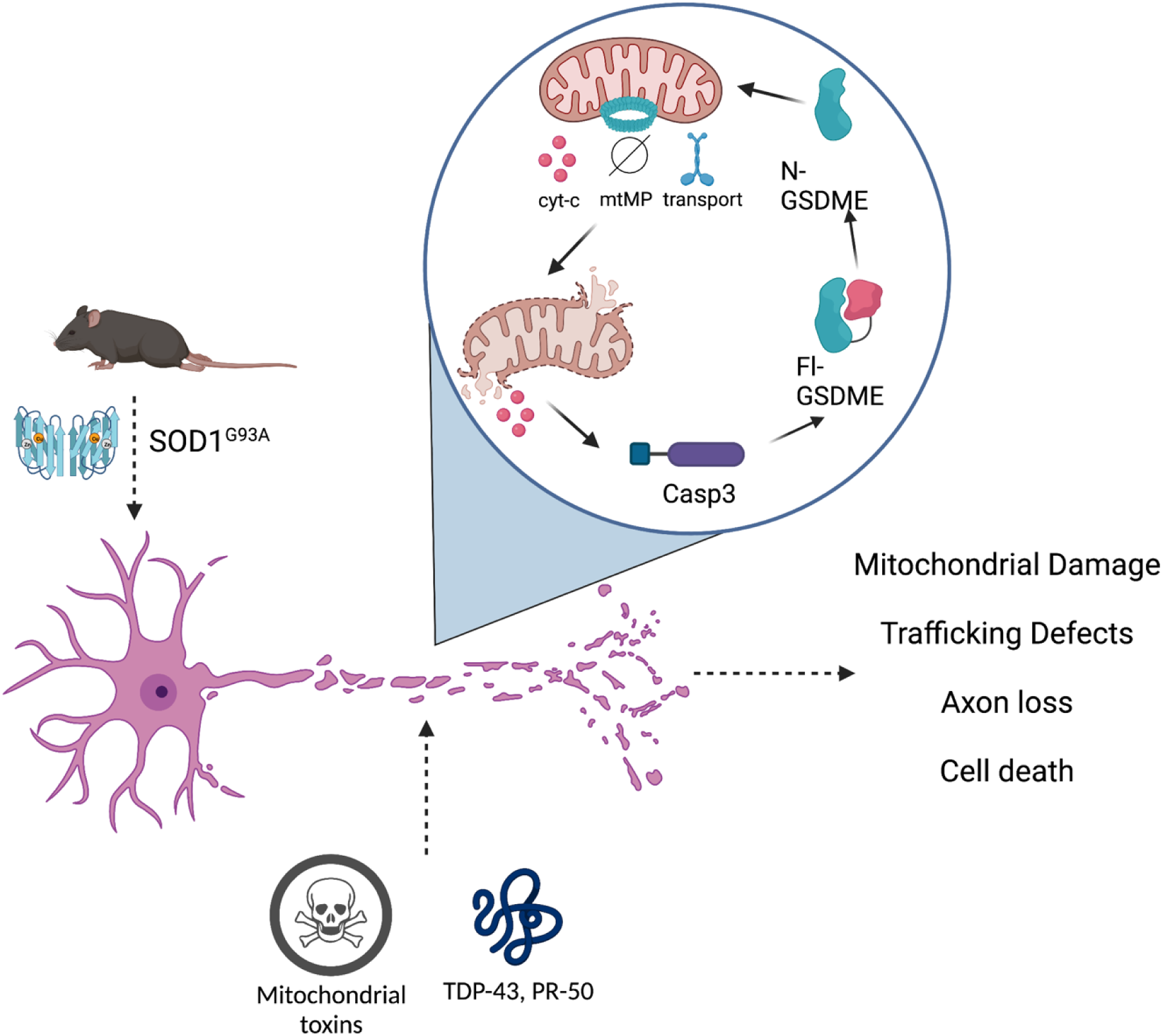

**Highlights:**

- GSDME is expressed by neurons and activated by mitochondrial neurotoxins
- Activated GSDME drives axonal mitochondrial damage and neurite loss prior to cell death
- ALS/FTD associated TDP-43 and PR-50 induces GSDME-driven neurite loss in mouse and human iPSC-derived neurons.
- SOD1^G93A^ mice show ameliorated disease progression and motor neuron loss in absence of GSDME

## Introduction

Neurologic diseases are often characterized by early mitochondrial dysfunction and axon loss, which precede overt cell death ^1–4^. Healthy neuronal mitochondria are essential for a variety of cellular functions, including efficient ATP generation^5^, synaptic transmission ^6^, ROS generation and calcium signaling ^6–8^. Mitochondrial damage leads to release of contents including mitochondrial DNA and cytochrome-c, which drives caspase-3 activation and apoptosis ^9^. Localized caspase-3 activity can also occur in neurites and contributes to axonal damage; once a threshold is reached in the soma, caspase-3 can drive overt neuronal cell death ^3^. Though the link between mitochondrial damage and caspase-3 activation is well characterized, the myriad mechanisms that amplify local mitochondrial demise in neurites, drive axonopathy and lead to neuronal dysfunction are not fully understood. Mapping the molecular mechanisms linking mitochondrial stress to axon degeneration and neuronal death may have implications for a wide range of neurologic insults.

Pyroptosis is a necrotic form of inflammatory cell death mediated by members of the pore-forming Gasdermin superfamily: GSDMA, GSDMB, GSDMC, GSDMD, and GSDME (or DFNA5). Gasdermins have pore-forming N-terminal domains that are masked by an autoinhibitory C-terminus in the resting state. Proteolytic cleavage in the linker region between the -N and -C domains release the N-terminal fragment of GSDMs to insert and oligomerize in lipid membranes and form pores. Accumulation of GSDM pores in the plasma membrane leads to cellular swelling and overt necrosis—a process termed “pyroptosis”^10–13^. To date, most studies concerning GSDM-biology have focused on immune, epithelial or cancer cells. GSDMD, which was first identified in macrophages, is enriched in immune cells, and activated by cleaved caspase 1/11 downstream of inflammasome assembly ^12–14^. Prior to mediating pyroptosis, N-GSDMD can also permeabilize mitochondrial membranes leading to release of cytochrome-c, mtDNA and downstream caspase-3 activation^15, 16^. However, the range of cell types where GSDM family members are expressed and activated has not been well defined.

In this study, we focus on the role of one family member, GSDME, based on its expression in brain and spinal cord. Prior to its discovery as a gasdermin family member, GSDME was first named deafness associated family member 5 (DFNA5). Patients with gain of function mutations in GSDME present with progressive sensorineural hearing loss—the hair cells of the cochlea are destroyed but other tissues appear to be spared^17, 18^. Early Northern blot and subsequent RT-qPCR analyses showed that DFNA5 was broadly present in mesenchymal-derived tissues, placenta, and bulk brain samples ^17, 19^. The molecular mechanism of GSDME activation was first elucidated in cancer cell lines, where it was shown to be cleaved by caspase-3^11^. Once released, the GSDME N-terminal fragment inserts into the plasma membrane and forms pores that drive pyroptosis ^10, 11^. Subsequent work has shown that GSDME can be activated in human keratinocytes ^20^, the GI tract ^21^, kidney ^22^, mouse retinoblastoma cells ^23^, and human cancer cells ^11, 24, 25^.

The function of GSDMs in neurons and its action in axonal biology has not been studied. Here, we find that both mouse and human neurons express GSDME, and that it is enriched in neurons compared to glia. Neurons are distinct from other GSDM-positive cell types; they are non-dividing and possess long morphological processes, namely dendrites and axons. Neurons also have unique energetic demands, needing to transport mitochondria over great distances to maintain signal transduction in axons and synapses. In neurodegenerative conditions such as Parkinson’s Disease (PD), frontotemporal dementia (FTD), and amyotrophic lateral sclerosis (ALS), proteins such as α-synuclein, TDP-43, SOD1 and *C9ORF72*-associated dipeptide repeat proteins (DPRs) disrupt mitochondrial function and induce loss of axonal processes. These changes are early steps in disease pathogenesis^1, 5, 26–29^.

Our study reveals that GSDME mediates local mitochondrial damage in axons that drives neurite loss and cell death. Activated GSDME rapidly distributes to neurite-associated mitochondria to amplify membrane potential loss, slow mitochondrial trafficking, and promote neurite retraction. In microfluidic chambers, we show that GSDME-driven mitochondrial damage can be localized to distal axons. FTD/ALS-associated proteins induced GSDME activation and localization to axonal mitochondria, driving mitochondrial depolarization and neurite loss. GSDME is activated and plays a functional role in mouse (SOD1^G93A^) and a patient-derived iPSC-motor neuron model of ALS. Genetic knockout or shRNA-based knockdown of GSDME protects against axonal damage to neurons *in vitro* and motor dysfunction *in vivo*. Thus, we identify GSDME as an important modifier of mitochondrial damage neuronal dysfunction that may play a role in neurologic disease.

## Results

### GSDME is expressed in mouse and human neurons

The expression of the Gasdermin family in the central nervous system and its potential functions in neurons have not yet been well characterized. We mined publicly available single cell RNA-seq data of mouse cortex and hippocampus sorted for neurons^30, 31^.The GSDM family members *Gsdmd*, *Gsdma*, and *Gsdmc* were largely absent, while *Gsdme* was expressed in both GABAergic and Glutamatergic neurons (**Fig 1A**) To further confirm *Gsdme* expression in the mouse brain, we isolated tissue samples from different brain regions of wild-type (WT) or *Gsdme*^-/-^ mice for transcriptional analysis, immunostaining, and immunoblotting. *Gsdme* mRNA was detected in cortical, cerebellar, midbrain and spinal cord lysates from WT mice, but absent in Gsdme^-/-^ (KO) animals (**Fig S1A**). Anti-GSDME immunostaining of mouse brain showed a broad expression pattern in cells with neuronal morphology (**Fig 1B**). As expected, staining and immunoblotting of mouse brains from knockout animals showed minimal immunoreactivity (**Fig S1B** and **Fig 1C**).

**Figure 1:**
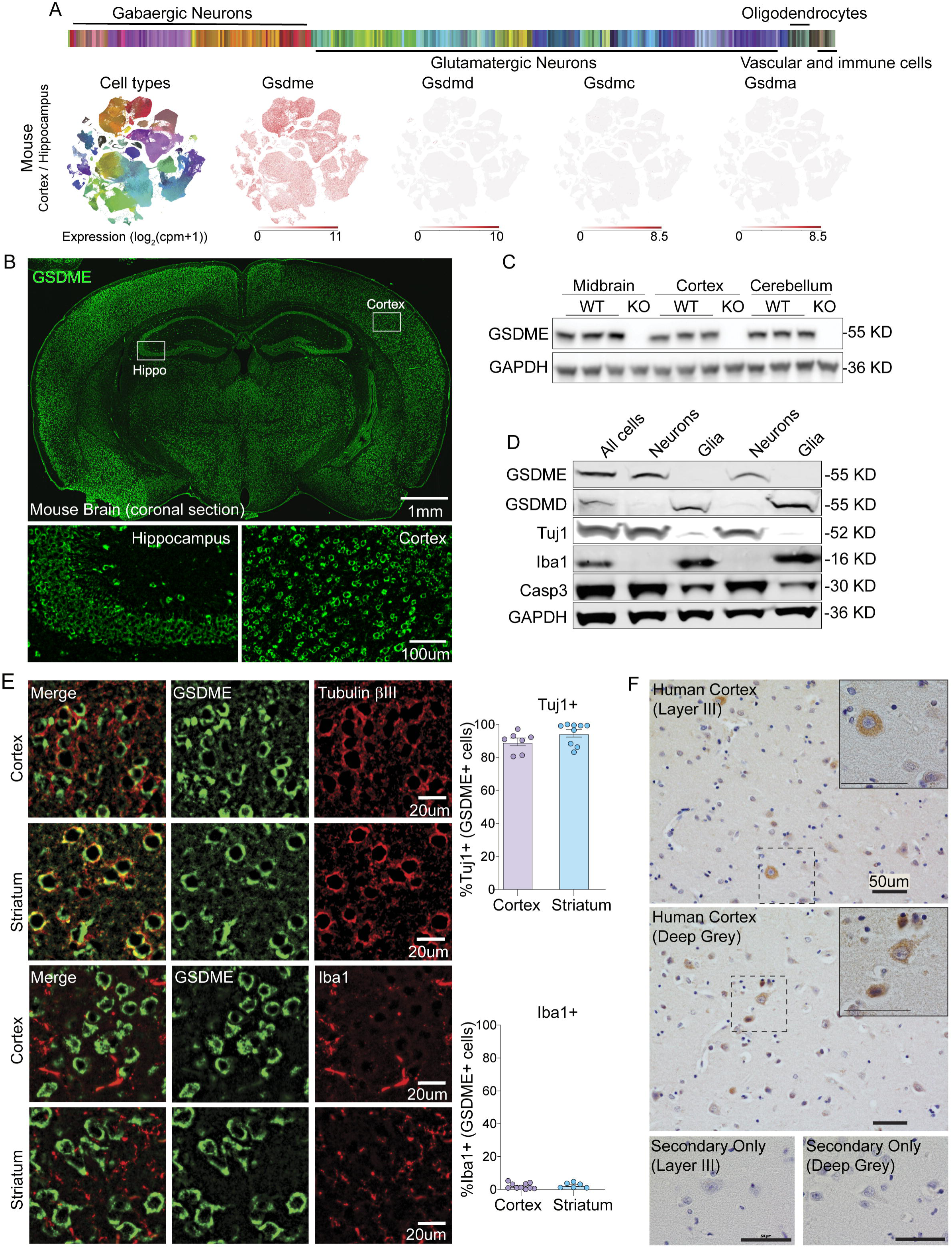
GSDME is expressed in the brain and localizes to neurons at baseline. (A) Publicly available single-cell RNA-seq (10X Genomics) data for mouse cortex and hippocampus enriched for neurons was mined for expression levels of Gasdermins. TSNE plots showing cell types (left most plot) and gasdermin genes were generated using the Allen Brain Atlas Transcriptomics Explorer tool (Allen Institute for Brain Science). (B) Representative IHC images of coronal mouse brain sections stained using anti-GSDME antibody. Scale bar: 1mm (C) Immunoblot analysis of GSDME expression in several mouse brain regions from adult wild-type and GSDME knockout mice. (D) Immunoblot analysis of full-length GSDME and GSDMD expression in Miltenyi-MACS enriched neuronal (Tuj1+) and glial (Iba1+) cell populations from P0 mouse pups. (E) Representative IHC images of mouse brain sections from cortex and striatum co-stained with anti-GSDME and either anti-beta-III-tubulin (Tuj1) or anti-Iba1 antibodies. Scale bar: 20um Colocalization analysis of GSDME positive cells with markers for neuronal (Tuj1) or glial (Iba1) cells. Results are displayed as the percentage of GSDME + cells that also express either Tuj1 or Iba1. Each dot represents a coronal mouse brain section. Two coronal sections were quantified per mouse, and n= 3-5 mice were used for each co-staining experiment. (F) Representative IHC images of temporal lobe from two healthy control patients stained with human anti-GSDME. Secondary only controls are displayed in smaller panels underneath the stained sections. Scale bar: 50um

To test if GSDME is enriched in neuronal cells compared to glia, we performed MACS purification of neurons vs. non-neuronal cells from mouse brain followed by western blot analysis (**Fig 1D**). We found clear GSDME signal in neuronal fractions that were also positive for neuronal beta-III-tubulin (Tuj1). GSDMD, which has been mostly characterized in macrophages ^14^, was absent in neurons. However, the non-neuronal fractions (Iba1+, Tuj1-) showed clear expression of GSDMD but not GSDME (**Fig 1D**). Consistent with this analysis, transcriptomic data ^32^ from FACS-isolated mouse microglia showed *Gsdmd* expression at baseline but sparse *Gsdme* detection in these same populations (**Fig S1C**). Co-staining sections of cortex and striatum showed high colocalization of GSDME with the neuronal marker Tuj1, and not the microglial marker Iba1 (**Fig 1E**), confirming neuron-specific enrichment of GSDME compared to microglia. Evidence from publicly available human single cell transcriptomic databases^30^.(Bakken et al., 2021) similarly demonstrated that *GSDME*, but not family members *GSDMD*, *GSDMA* or *GSDMC,* is expressed at baseline in human neurons (**Fig S1D**). We next analyzed GSDME protein expression in human brain samples. Temporal cortices from human brains were stained with anti-GSDME. We found that GSDME was expressed in multiple cortical layers; these GSDME-positive cells showed a perikaryal staining pattern consistent with cytosolic expression in neurons (**Fig 1F**). Taken together, this data shows that GSDME is expressed at baseline in both mouse and human neurons.

### GSDME mediates neuronal cell death caused by mitochondrial toxins

Mitochondrial toxins, such as rotenone and 6-OHDA, are intrinsic apoptotic stimuli often used to induce neuronal cell death, and model features of neurodegeneration *in vitro* and *in vivo*^2, 4^. In cancer cell lines, GSDME is classically cleaved and activated by caspase-3 downstream of apoptotic triggers (**Fig 2A**)^11^. Given that this molecule is expressed in neurons, we sought to understand if toxin treatment could activate GSDME in human and murine cells. For this study we utilized several mitochondrial toxins: rotenone and 6-OHDA (complex I inhibitor), antimycin (complex III inhibitor) and raptinal (a rapid inducer of mitochondrial damage and caspase-3). We found that treatment of primary mouse cortical neurons and human SH-5YSY cells with these mitochondrial toxins led to cleavage of full-length GSDME to the ∼30 kDa N-terminal pore-forming fragment by immunoblot analysis (**Fig 2B & S2A**) and cell death as measured by lactate dehydrogenase (LDH) release and uptake of the membrane impermeant dye propidium iodide (PI) (**Fig S1E-H**). We confirmed specific GSDME cleavage following toxin treatment through immunoblot analysis of WT and GSDME KO SH-SY5Y cells generated using CRISPR-Cas9 targeting (**Fig S2A**). Thus, these toxins can be used as probes to explore the biological functions of neuronal GSDME.

**Figure 2:**
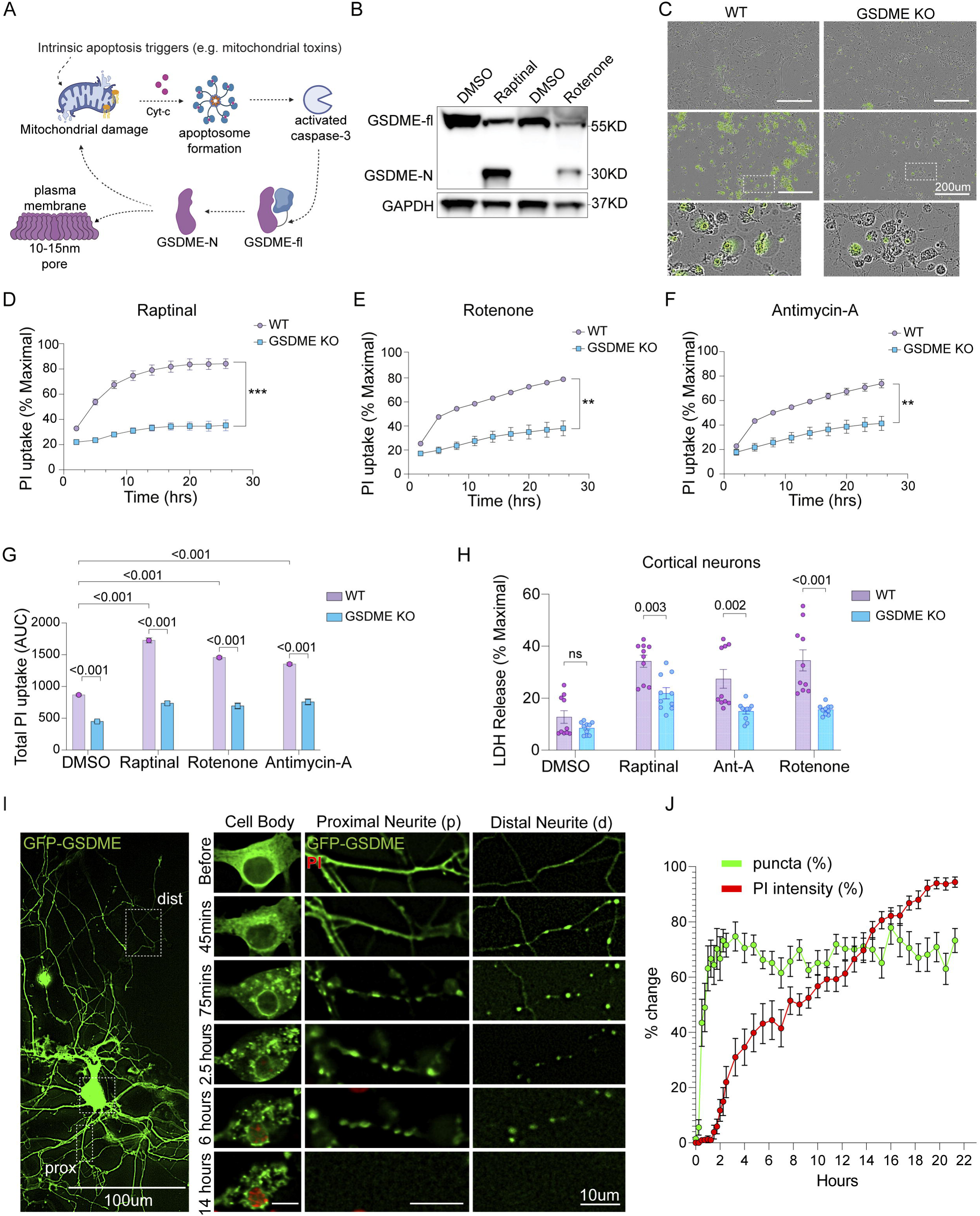
GSDME is activated by mitochondrial toxins and contributes to cellular necrosis in neurons. (A) Schematic of GSDME activation downstream of intrinsic apoptotic stimuli and caspase-3 processing. Yellow lipids on the outer mitochondrial membrane represent cardiolipin residues. (B) Immunoblots of primary neurons treated with 10uM of raptinal for 1h or 20uM rotenone for 8h. (C) Representative 20X Images of WT (top panels) or GSDME KO (bottom panels) primary mouse cortical neurons at 3h following treatment with 5uM raptinal or DMSO (Control). Cells were incubated in *Sytox* Green-containing media, and green cells indicate dye uptake. Scale bar: 200 um (D-F) Primary neurons were incubated in propidium iodide (PI) containing media and treated with (D) 4uM raptinal, (E) 20uM rotenone or (F) 30uM Antimycin-A. Images were taken every 3h for a span of 24 hours and quantified for PI uptake. (G) Area under the curve (AUC_24h_) measurements for cells treated with toxins as indicated in panels D-F (H) WT and GSDME KO mouse cortical neurons were treated with 4uM raptinal (1h), 20uM rotenone (2h) or 20uM Antimycin-A (1h) were assessed for LDH release at 24h post-treatment. (I-J) Representative images of primary neurons transfected with GFP-GSDME and incubated in media containing propidium iodide (PI). These cells were treated with 5uM raptinal and imaged over the course of 21h. The left (large) panel shows a single transfected neuron prior to any drug treatment. Distal neurites (dist) were defined as segments >200 um from the cell body, while proximal neurites (prox) were located < 200 um away. Images were captured every 15 min and (J) the formation of intracellular puncta, as well as PI uptake were quantified over time in the neuronal cell bodies. Each data point represents an average of 15 neurons. Data represents an average of at least 2-3 independent experiments ± SEM. Each independent experiment had 4 technical replicates (wells in 96 well plate) per condition. For datasets (D-F) two-way ANOVA (row factor = toxin treatment, column factor = genotype) followed by multiple comparisons was done to compare each group (adj p-values were calculated by the Tukey method).

We next determined whether loss of *Gsdme* impacts cell death downstream of mitochondrial toxin treatment. *Gsdme* KO primary mouse neurons showed significantly reduced PI or Sytox Green uptake compared to WT neurons treated with the toxins raptinal, rotenone and antimycin-A (Ant-A) (**Fig 2C-G**). These results were confirmed in GSDME KO SH-SY5Y cells, which were also protected from PI uptake induced by raptinal, rotenone, 6-OHDA, Ant-A and 3-NP, suggesting that GSDME mediates cell death downstream of multiple mitochondrial toxins (**Fig S2B-G)**. Challenge with ER stressors tunicamycin and thapsiagargin, which generate ROS and activate caspase-3, also caused a GSDME-dependent PI uptake (**Fig S2I-K**). These findings indicate that GSDME contributes to plasma membrane permeability both in primary mouse neurons and a human neuron-like cell line, downstream of multiple neurotoxic triggers. Toxin-induced LDH release was also partially rescued in GSDME deficient primary neurons and SH-SY5Y cells compared to wild-type controls (**Fig 2H & S2H**). Taken together, these experiments show that mitochondrial toxin induced necrosis (PI+ uptake and LDH release) is regulated by GSDME in human neuroblastoma cells and mouse primary cortical neurons.

### GSDME activation in neurons is caspase-dependent

Given that mitochondrial toxins classically activate caspase-3, which is upstream of GSDME cleavage^10, 11, 20^, we next assessed whether GSDME-driven cell death is indeed caspase-dependent in neurons. Treatment with the pan-caspase inhibitor zVAD-FMK blocked raptinal-induced GSDME processing into its N-terminal fragment (**Fig S3A-B**). Incubation of SH-SY5Y and mouse primary cortical neurons with zVAD-FMK phenocopied the effect of GSDME knockout, decreasing PI uptake after toxin treatment (**Fig S3C-F**). By contrast, treatment with the RIP kinase inhibitors nec-1s and GSK ‘872 did not reduce toxin induced PI uptake in primary neurons, suggesting that necroptosis does not participate in mitochondrial toxin-driven neuronal death (**Fig S3G-H).** These data indicate that treatment of neurons with mitochondrial toxins activates a caspase-3-GSDME axis, leading to neuronal death.

### Activated GSDME rapidly localizes to mitochondria

Activated (N-terminal) GSDME can bind to a range of acidic lipid residues, such as phosphatidyl serine in the plasma membrane and cardiolipin (CL) in mitochondria.^11, 16, 34^ Following insertion into lipid bilayers, N-GSDM fragments oligomerize to form 10-21nm pores.^11, 13^ Neurons have large somas and fine dendritic and axonal processes (neurites). Given the unique morphology and cell biology of neurons, we hypothesized that activated GSDME may act at specific subcellular compartments to drive cellular dysfunction. To investigate the intracellular dynamics of GSDME, before and after activation, we transfected mouse cortical neurons with N-terminally tagged full-length GFP-GSDME (**Fig S4A**). We validated that GFP-GSDME maintained pore-forming and pyroptotic ability by transfecting this construct into HEK293T cells, which lack endogenous GSDME **(Fig S4B-C)**. When transfected cells were treated with raptinal, we observed caspase-3 activation, cleavage of GFP-GSDME, rapid PI+ uptake and pyroptotic morphology **(Fig S4C-F)**. These changes were not observed in untransfected or GFP-transfected control cells, which lack GSDME expression **(Fig S4D)**. Performing immunoblot analysis of the time course for GSDME activation, we observed that cleavage of both endogenous GSDME and transduced GFP-GSDME were identical and concomitant with caspase-3 cleavage in primary neurons (**Fig S4G**).

Following raptinal treatment of neurons, GFP-GSDME rapidly formed numerous intracellular puncta in the distal neurites by 45 minutes, and eventually proximal neurites and cell bodies by 75 minutes (**Fig. 2I-J**). When cortical neurons were co-transfected with a membrane-RFP marker and treated with raptinal, GFP-GSDME puncta appeared predominantly dispersed in the cytosol, with little or no colocalization with the plasma membrane (**Fig S5C**). This stands in contrast with findings in the SH-SY5Y cells, which had both plasma membrane and cytosolic enrichment of GFP-GSDME (**Fig S5A & Supp video 1**). The appearance of GSDME puncta in the distal and proximal neurites preceded PI uptake, indicating cell death, by several hours (**Fig 2I-J**). To assess whether the neurite associated puncta represent local sites of activated GSDME, we pre-treated the neurons with zVAD-FMK (caspase inhibitor), at a concentration that blocked GSDME cleavage **(Fig S3A-B).** Early toxin-induced puncta formation was inhibited by zVAD-FMK (**Fig S5A, S5D-E & Supp video 1**. This data suggests that in neurons, caspase dependent GSDME localization may occur earlier and to a larger extent to intracellular membranes than to the plasma membrane.

Previous work has shown that mitochondrial toxins such as rotenone, 6-OHDA and others cause cardiolipin to be externalized to the OMM in both primary cortical neurons and SH-SY5Y cells^33, 34^. Given the ability of GSDME to bind CL^11^, we next tested the hypothesis that GSDME may be localized to mitochondria along the neurites. In live cell imaging studies of cortical neurons, cells were co-transfected with GFP-GSDME and with mCherry-OMP25 to allow visualization of the outer mitochondrial membrane (OMM). We analyzed the fraction of GSDME that colocalized with OMP25 in the distal axons and found raptinal treatment resulted in a 1.5-fold increase enrichment of GSDME on axonal mitochondria and soma (**Fig. 3A-C and S6A-B**). In addition to raptinal, the mitochondrial toxin rotenone also caused rapid co-localization of GFP-GSDME with mCherry-OMP25+ mitochondria (**Fig S6C-E**). Immunoblot analysis of fractionated lysates from raptinal-treated SH-SY5Y cells (**Fig S6F**) and primary cortical neurons revealed a three-fold enrichment endogenous cleaved GSDME in mitochondria compared to cytosolic fractions (**Fig S6G-J**). GSDME colocalization with mitochondria was also associated with release of the intermembrane/intercristae space protein, cytochrome-c, into the cytosol (**Fig S6 I, K-L**). To validate the cardiolipin-dependent docking of GFP-GSDME on mitochondria we knocked down cardiolipin-synthase (CLS-1) in primary neurons. CLS-1 knocked down neurons showed decreased GFP-GSDME puncta formation post toxin treatment (**Fig S7 A-C**). This data suggests that following activation, GSDME localizes to neuronal mitochondria in a cardiolipin-dependent manner.

**Figure 3:**
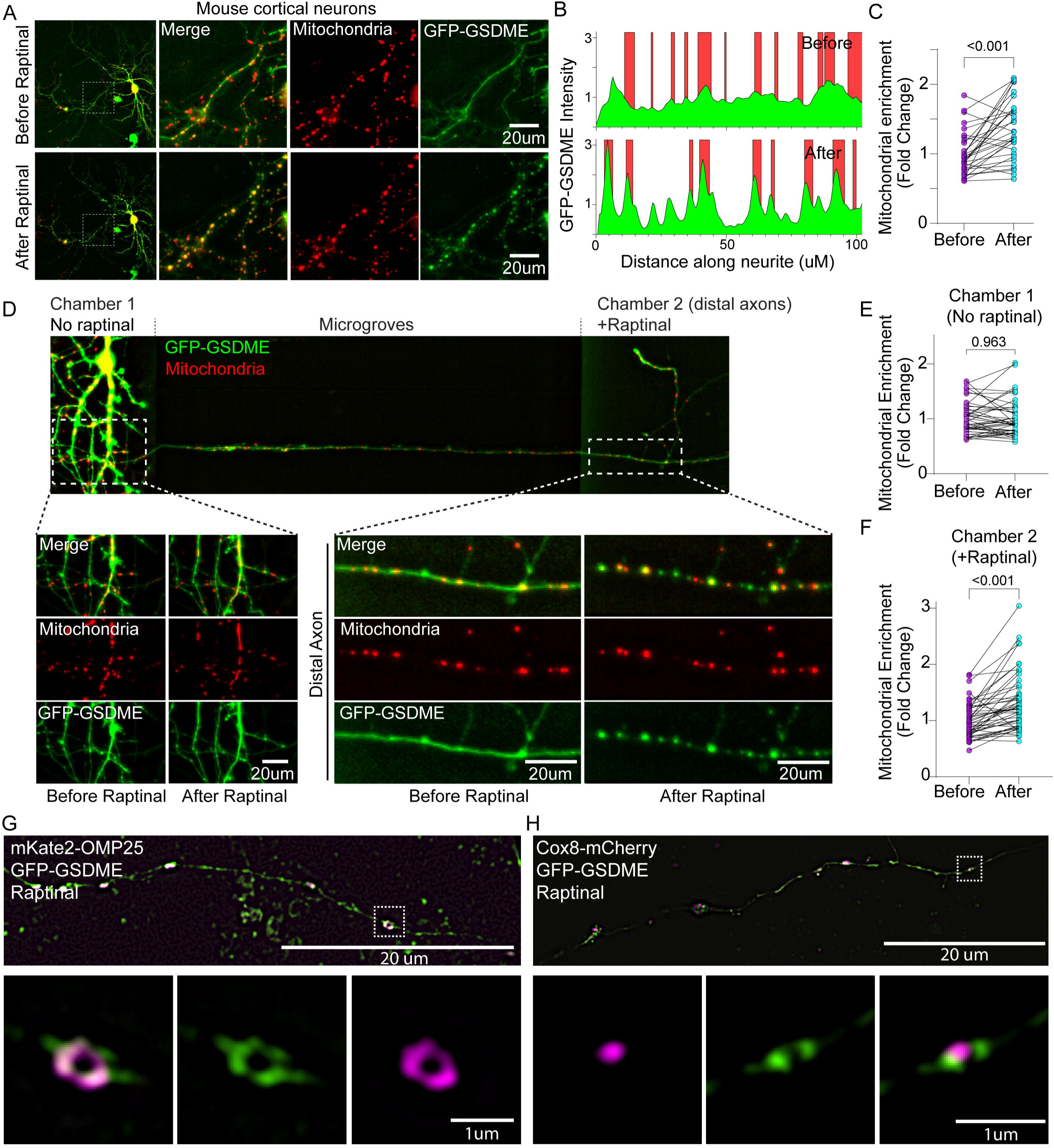
Activated GSDME rapidly localizes to neuronal mitochondria following toxin treatment. (A-C) Mouse neurons co-transfected with mitochondrial marker mKate-OMP25 and GFP-GSDME were imaged before and after 90mins of treatment with 5uM raptinal. Representative images of a neuron before and after treatment is shown (left panels). Middle and right panels show a magnified image of a neurite (indicated by white dotted box), with the red (mitochondrial) and green (GSDME) signals separated. (B) The location of mitochondria (red) and GFP-GSDME (green) peaks as measured via line scans along the neurites are indicated. (C) The enrichment of green GSDME signal at mitochondria (red) over the background (diffuse cytosolic intensity) were quantified from line scans along neurites (as shown in panel B) and compared before and after raptinal treatment. N= 30 neurite segments representing neurons from 3 biological replicates. (D-F) Representative image of a microfluidic chamber plated with wild-type mouse cortical neurons transfected with GFP-GSDME and mKate-OMP25 is shown. The cell body, microgrooves and axonal compartments are labeled (D). The axonal chamber was treated with 2uM raptinal for 2.5h. Images of proximal neurites (left) and a distal axon segment (right) before and after raptinal treatment are shown. Enrichment of GFP-GSDME on mitochondria in (E) proximal chamber 1 neurites and (F) distal chamber 2 axons before and after raptinal treatment were quantified (N = 50 distal axon segments and 50 proximal neurite segment representing neurons from three separate microfluidic chambers.) (G-H) Representative images of mouse neurons transfected with GFP-GSDME and either (G) OMP25-mCherry or (H) Cox8-mCherry. These cells were treated with 10uM raptinal were fixed and imaged using structured illumination microscopy (SIM) to assess marker colocalization. For datasets C, E, and F paired Student’s pairwise t-tests were performed to compare neurons before and after drug treatment, Each dot represents a neurite segment. The combined datasets represent individual neurons taken from 2 or 3 independent cell culture experiments.

We then asked whether local toxin-exposure can induce GSDME activation in axons, without involvement of the soma. We plated mouse cortical neurons in microfluidic chambers, which allowed for selective treatment of the axonal compartments with toxin (**Fig 3D**). We found that raptinal caused GFP-GSDME puncta formation and mitochondrial colocalization that was specific for the axonal compartment and did not occur in the untreated soma or proximal neurites of the same cell (**Fig 3D-F**). These data suggest that mitochondrial stress leads to enrichment of GSDME on neuronal mitochondria and can occur locally in axons.

We next used high resolution structured illumination microscopy (SIM) of neurons transfected with full-length GFP-GSDME and treated with raptinal. This analysis revealed robust puncta formation that colocalized with the outer mitochondrial membrane (OMM) marker mKate-OMP-25, but not with the matrix marker Cox8-mCherry (**Fig 3G-H; Supp Videos 2-3**). These results suggest that toxin treatment targets activated GSDME to mitochondrial membranes, and not the matrix.

### Neurotoxins induce GSDME cleavage and mitochondrial localization in human iNeurons

Human iPSC derived neurons (iNeurons) are postmitotic, electrically active, and thus more representative of human neurons than SH-SY5Y neuroblastoma cells^35^. Here, we generated glutamatergic neurons that are most similar to layer 2/3 glutamatergic neurons of the cerebral cortex^36^. We investigated whether GSDME is similarly activated in these cells. We found raptinal treatment induces a dose dependent GSDME cleavage in iNeurons (**Fig S8A**). The activation of GSDME in iNeurons was blocked in the presence of the caspase inhibitor zVAD-FMK (**Fig S8B).** Human iNeurons transfected with GFP-GSDME and mCherry-OMP25 also showed robust mitochondrial colocalization upon toxin treatment, with a large number in the distal neurites (**Fig S8C-E**).

### Gsdme deficiency protects neurons from mitochondrial damage

We next asked whether GSDME enrichment on neuronal mitochondria potentiates damage to these organelles. To assess the functional consequences of GSDME mitochondrial colocalization in neurons, we performed TMRM staining following toxin treatment. TMRM is a voltage sensitive dye sequestered by healthy, polarized mitochondria. TMRM rapidly loses fluorescence following mitochondrial damage and membrane depolarization^37^. Treatment of wild-type primary neurons with raptinal, the complex I inhibitor rotenone, or the complex II inhibitor antimycin-A, leads to dose dependent mitochondrial depolarization and loss of TMRM stain. Depolarization was partially rescued in *Gsdme* KO primary cortical mouse neurons and SH-SY5Y (**Fig 4A** and **Fig S9A-D**). Though the timing and extent of rescue differ between cortical neurons and SH-SY5Y cells, toxin-treated GSDME KO cells consistently had higher TMRM staining than WT counterparts. These experiments suggest that downstream of an initial mitochondrial insult (i.e., toxin exposure), GSDME potentiates or accelerates mitochondrial damage. To assess how GSDME impacts mitochondrial membrane integrity we transfected WT and KO cortical neurons with mKate2-OMP25 (OMM protein) and treated with toxin. After 2h and 4h of treatment (but not by 1h), Gsdme KO neurons had preserved OMP-25 signal, suggesting a sparing of the OMM relative to WT cells (**Fig 4B-C**). To directly visualize the structural integrity of mitochondria, we performed transmission electron microscopy (TEM) of WT and *Gsdme* KO neurons treated for 1h with raptinal. Analysis of TEM images revealed that raptinal-treated WT mitochondria had less pronounced cristae and increased fragmentation (reduced length) compared to those in *Gsdme* KO neurons (**Fig 4D-E**). Raptinal-treated WT neurons also displayed a high percentage of mitochondria with damaged inner or outer membranes, while *Gsdme* KO neurons were almost completely rescued (**Fig 4F**). This suggests that GSDME increases mitochondrial membrane damage downstream of toxin-mediated electron transport chain disruption.

**Figure 4:**
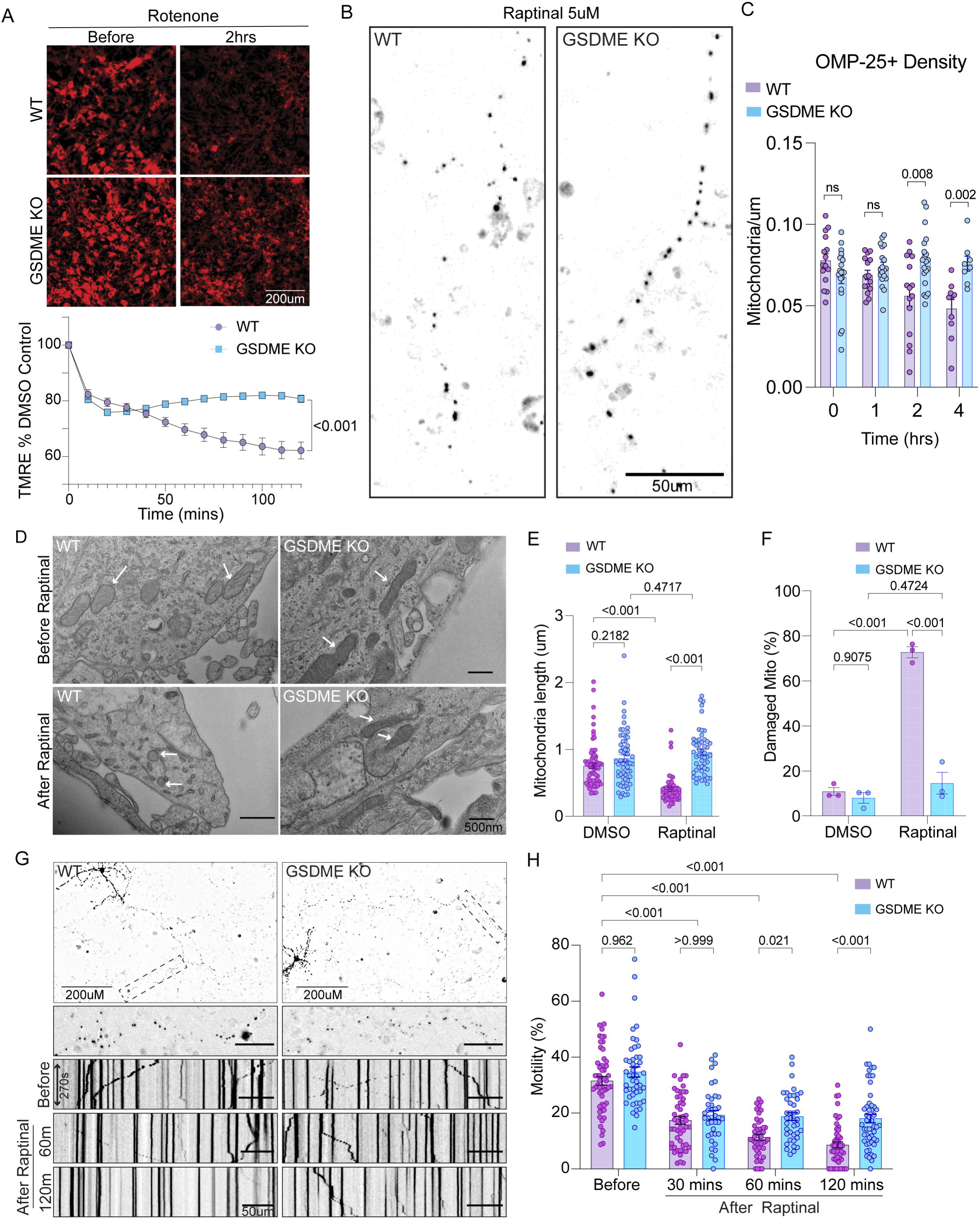
GSDME deficient neurons are protected from toxin-induced mitochondrial dysfunction. (A-) Wild-type and GSDME knockout primary neurons were stained with TMRM and incubated with 20uM rotenone Images were captured every 15 minutes and TMRM intensity normalized to DMSO controls was quantified. (B-C) Wild-type and GSDME knockout primary neurons were transfected with OMP25-mCherry and then treated with 5uM raptinal. (B) Representative images of cells imaged at 0, 1h, 2h and 4h post-raptinal treatment. (C) Mitochondrial density was calculated by counting the number of OMP-25+ objects and dividing by neurite length (D-F) Representative transmission electron microscopy images (TEM) of WT and GSDME KO neuronal cultures treated with either DMSO or raptinal for 1h, and then fixed, dissociated and processed for TEM. Both the (E) mitochondrial length (each dot = 1 mito) and (F) percentage of damaged mitochondria (each dot = average of 2 wells) were calculated across three independent experiments. (G) Wild-type and GSDME KO mouse neurons were transfected with OMP25-mKate, treated with 3uM raptinal and imaged at high time resolution (1 image/5s) for 3 min intervals. These intervals were captured at 0h, 1h and 2h post-toxin exposure. Kymograph analysis was performed to visualize and quantify mitochondrial motility. (H) Percent motile mitochondria was calculated from kymograph analysis of WT and GSDME KO neurons treated with 3uM raptinal. Combined data from three independent experiments are shown. For datasets A-C, student’s t-tests were performed for comparisons between genotypes, and p-values were adjusted using the Tukey method. Each dataset represents an average of two independent experiments ± SEM. For datasets E and H two-way ANOVA was performed, followed by multiple comparisons for each group. Adjusted p-values are mentioned.

Cyt-c release into the cytosol indicates outer mitochondrial membrane (OMM) damage. Raptinal treatment caused rapid (<1h) cyt-c release in WT cortical neurons and SH-SY5Y (**Fig S6 G, I**). Immunoblots for cyt-c revealed that toxin treated GSDME KO SH-SY5Y displayed a significantly higher mitochondrial to cytosolic cyt-c ratio, relative to WT cells (**Fig S6 K-L**). This indicates that mitochondria from toxin-treated KO cells had less OMM damage (i.e., cyt-c release) compared to WT cells. Consistent with less mitochondrial depolarization and cyt-c release, GSDME KO SH-SY5Y also exhibited reduced caspase-3 activation relative to WT cells (**Fig S10A-C**). Though this feed forward amplification of caspase-3 by GSDME has been described in other cell types (**Fig 2A**)^38^, we document it in SH-SY5Y cells exposed to complex I inhibitors and PD-associated toxins (e.g., rotenone and 6-OHDA). *Gsdme* KO cortical neurons treated with raptinal also had partially reduced caspase-3 activation relative to WT neurons (**Fig S10D-E**) albeit to a lesser extent than the SH-SY5Y cells. When taken together, these data indicate that GSDME participates in a positive feedback loop to amplify mitochondrial damage and downstream caspase-3 activation in SH-SY5Y and primary mouse neurons.

Destruction of axonal mitochondria or disruption of trafficking to distal axons is an early step in many neurodegenerative diseases^1, 3^. If GSDME destroys mitochondrial membranes following toxin exposure, we reasoned that fewer mitochondrion would resume normal trafficking after toxin withdrawal. To determine the role of GSDME in impacting mitochondrial trafficking, we transfected primary neurons with mKate-OMP25 and took high timescale images in WT and *Gsdme* KO cells treated with toxins. At baseline in primary mouse neurons, we noticed no differences in mitochondrial movement (percent motile) between the two genotypes (**Fig 4G-H**). Treatment of WT cultures with the mitochondrial toxins raptinal and rotenone caused a striking arrest of motility at 1h and 2h post-treatment, as displayed in kymograph analyses (**Fig 4G**). This arrest was reduced in *Gsdme* KO cells, which had more motile mitochondria at these timepoints (**Fig 4H**).

### GSDME regulates neurite integrity and local mitochondrial potential

Axons require a constant supply of energy to maintain electrical excitability, structural integrity and synaptic function^1^. Given that GSDME promotes mitochondrial depolarization and reduces motility following toxin exposure, we next hypothesized that GSDME KO neurons would have reduced degeneration of distal axonal processes. When treated with toxins, WT neurons displayed a dose dependent increase in the ratio of depolymerized to polymerized beta III-tubulin (**Fig S10F-G**). Higher levels of depolymerized tubulin indicate microtubule disassembly and axonal degeneration^39^.

Following 30 min (raptinal, antimycin-A) or 2h (rotenone) treatment with toxins, cells were incubated in fresh media for 8h and then stained for beta-III-tubulin (Tuj1) (**Fig 5A**). Toxin-treated WT cells exhibit a higher ratio of depolymerized (clumpy and round) to total beta-III-tubulin; GSDME knockout cells demonstrated a striking preservation of axonal processes containing polymerized tubulin (**Fig 5B-D**). At these timepoints, release of LDH from WT and KO neurons was minimal, indicating that GSDME increases axonal destruction prior to overt cellular necrosis (**Fig S10H, compare with Fig 2H**). Incubation of toxin-treated WT cells with zVAD-FMK also reduced the microtubule depolymerization index, phenocopying the effect of GSDME knockout (**Fig S10I**). Caspase inhibition rescue of neurite retraction also accords with findings from prior studies^3, 40^. Collectively, these experiments suggest that the caspase-3/GSDME axis is necessary to disrupt mitochondrial health and motility and to drive neurite loss before it contributes to overt cell death at the soma (i.e. PI uptake, LDH release).

**Figure 5:**
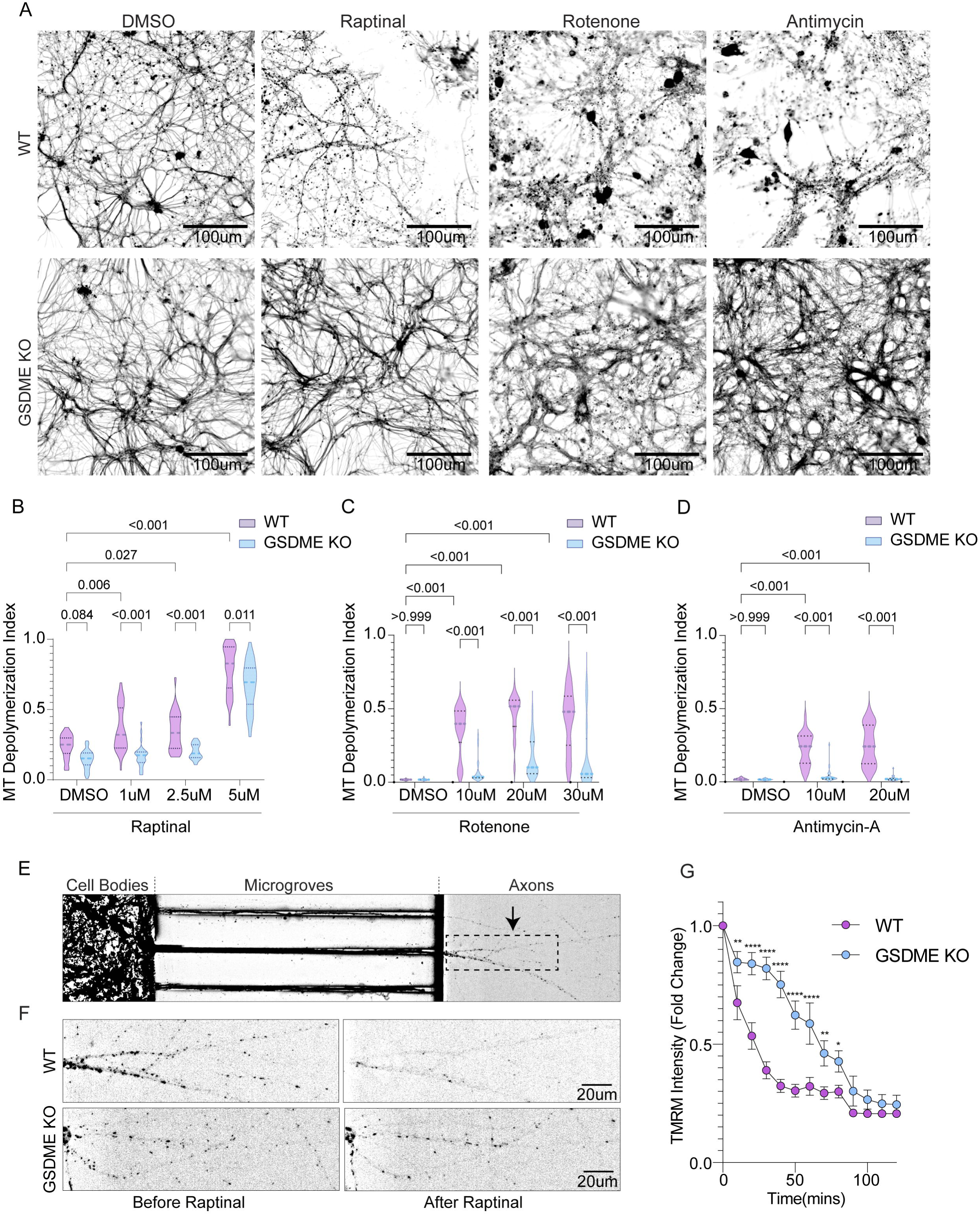
GSDME knockout protects against neurite loss and local mitochondrial damage. (A) Representative images of wild-type and GSDME knockout neurons treated with DMSO, 5uM raptinal, 20uM rotenone and 10uM antimycin-A, and stained for Tuj1 at 8h post-toxin treatment. (B-D) Microtubule depolymerization index was calculated for WT and KO neurons treated with several doses of (B) raptinal (C) rotenone or (D) antimycin-A. Violin plots display the combined median and interquartile ranges for depolymerization index taken from three independent experiments. (E) Representative image of a microfluidic chamber plated with wild-type mouse cortical neurons stained with TMRM. The cell body, microgrooves and axonal compartments are labeled. The panel represents the chamber prior to addition of 5uM raptinal. The black dashed box indicates an axonal region magnified in the following panel. (F) Representative images of a matched axonal segments from WT and GSDME KO neurons before (0 min) and after (60 min) addition of 5uM raptinal. Black signal indicates polarized (healthy) mitochondria stained with TMRM, and loss of signal indicates mitochondrial depolarization. The top panels show an axonal segment from WT neurons treated with raptinal, while the bottom panels show the effect of raptinal on GSDME KO neurons (G) Quantification of TMRM intensity relative to baseline (time = 0) from the axonal chambers of plated WT and GSDME KO neurons. Axonal segments from three WT and three KO microfluidic chambers were used for data analysis. This dataset represents the average of three independent experiments/chambers (n=3). For datasets B-D,E, two-way ANOVA (row factor = toxin dose, column factor = genotype) was performed, followed by multiple comparisons for each group (adj p-values, Tukey method).

We next asked whether the N-terminal GSDME fragment (N-GSDME) is sufficient to drive neuronal dysfunction. Expression of N-GSDME in primary mouse neurons caused localization to mitochondria in cell bodies and axons in the absence of any mitochondrial toxin treatment (**Fig S11A-B**). SIM microscopy analysis revealed overlap of GFP-N-GSDME and OMP-25 in both neurons and SH-SY5Y cells. However, only in SH-SY5Y cells did we observe GFP-GSDME enrichment on plasma membrane (**Fig S11B-D**). Expression of N-GSDME in primary cortical neurons was sufficient to cause mitochondrial potential loss (**Fig S11E-F**) and neurite damage as measured by Tuj1 staining (**Fig S9G-H**). Thus, N-GSDME expression phenocopies results obtained following chemical activation of GSDME using toxin treatment.

Using microfluidic chambers, we found that GSDME can be locally activated in axons and rapidly colocalizes with mitochondria (**Fig 3D**). To determine the effect of this local GSDME activation on axonal mitochondrial health, we again plated neurons in microfluidic chambers and stained with TMRM (**Fig 5E**). Following raptinal treatment of the axonal chamber, we observed dose-dependent TMRM decreases only in the treated axons (**Fig S10J-K**). Next, we examined WT and GSDME KO axonal compartments treated with toxins for differences in TMRM staining. Raptinal caused a rapid and pronounced loss of TMRM positivity, which was partially rescued in GSDME KO neurons (**Fig 5E-F**). While long term raptinal exposure leads to total TMRM loss, brief (30min-2h) treatment causes mitochondrial depolarization that is GSDME-dependent (**Fig 5G**). These data demonstrate that GSDME can be activated locally to destroy mitochondria, in the absence of overt changes at the cell body.

### ALS/FTD-associated proteins induce GSDME activation in neurons

Neurodegenerative conditions are characterized by early axonopathy that precedes loss of neuronal soma^26, 27^. Given that GSDME can drive axonal mitochondrial damage and disrupt structural integrity, we examined whether neurodegeneration-associated proteins would similarly activate GSDME. ALS and FTD are devastating neurodegenerative conditions characterized by progressive and ultimately fatal neuronal cell death. Both conditions are characterized by early mitochondrial damage in neurons and axon loss ^26, 41, 42^. We hypothesized that GSDME could be activated in these settings and potentially contribute to neuronal dysfunction.

To test whether ALS/FTD associated proteins cleaved GSDME in our *in vitro* system, we focused on the role of TDP-43 and the *C9ORF72* hexanucleotide expansion-associated dipeptide repeat protein (DPR) PR-50. TDP-43 is an RNA binding protein that forms neuronal inclusions in both familial and sporadic ALS/FTD brain and spinal cords^43^. Prior work has shown that TDP-43 aggregates bind mitochondria, leading to complex I inhibition and mitochondrial depolarization^44–46^. The DPRs produced from mutated *C9ORF72 have* also been shown to impair mitochondrial function (complex V activity), leading to caspase-3 dependent axon loss ^39, 47, 48^. Given that these neurodegenerative proteins are associated with direct mitochondrial damage and caspase-3 activation, we hypothesized that GSDME may be involved in downstream neuronal dysfunction mediated by these proteins.

Mouse cortical neurons were transfected with TDP-43, PR-50 or RFP control plasmids to model neurodegeneration in a dish ^39^.Co-expression of TDP-43 or PR-50 with GFP-GSDME, robustly increased GFP-GSDME puncta (**Fig 6A-B**). These puncta were enriched on neurite mitochondria, mirroring results obtained using complex I inhibitors. Treatment of WT cortical neurons with lentiviruses encoding control (GFP), TDP-43 or PR-50 under control of a Synapsin-I promoter, led to cleavage of endogenous GSDME at 72h post-transduction, indicating its activation (**Fig 6C** and **Fig S12A**).

**Figure 6:**
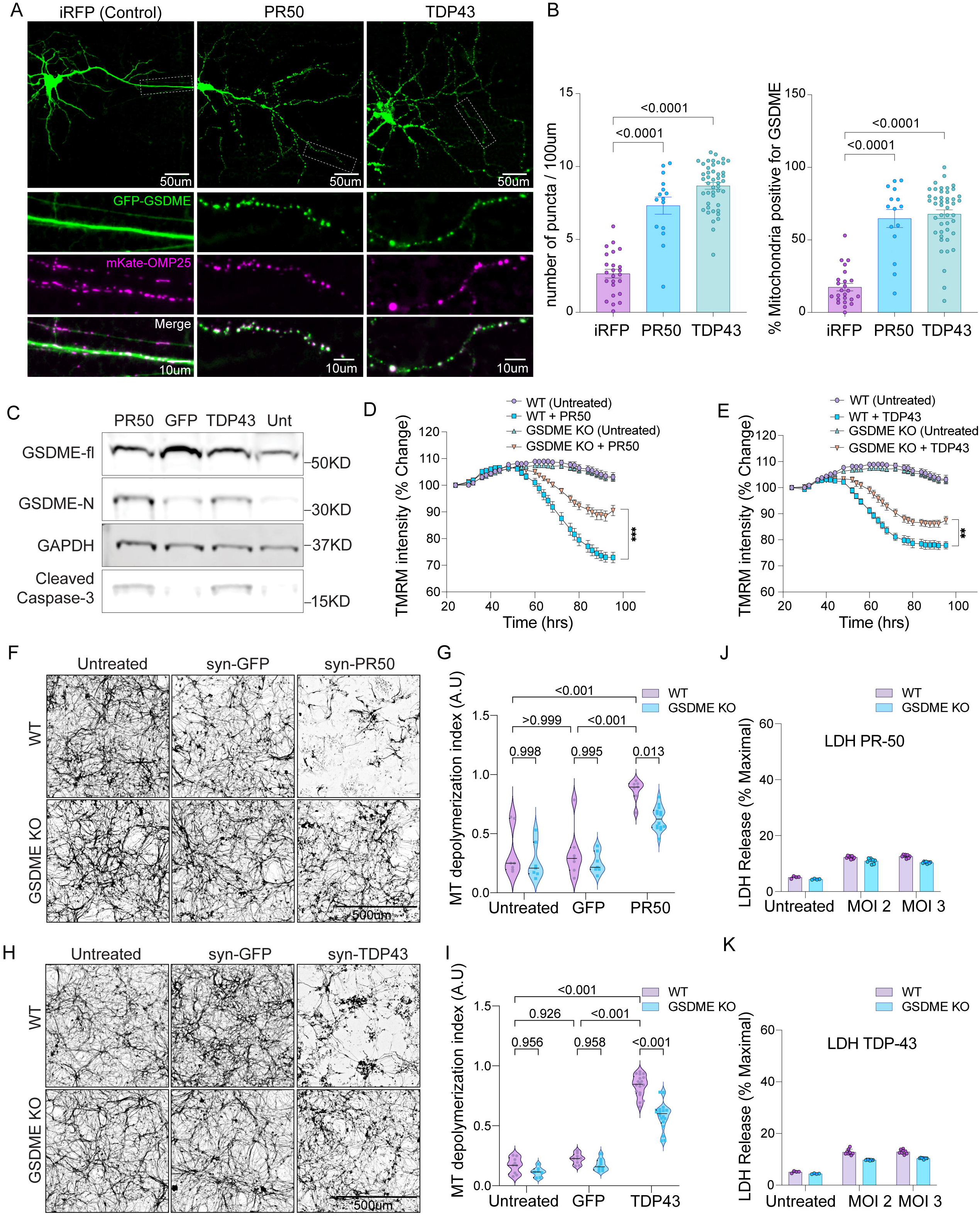
FTD/ALS proteins cause GSDME-dependent mitochondrial depolarization and neurite loss. (A) Primary mouse neurons were transfected with GFP-GSDME, mKate-OMP25 and either PR-50, TDP-43 or iRFP (control) constructs. Neurons were imaged at 48h post-transfection. The top panels show neurons in the GFP panel, and white boxes indicate magnified axonal segments. Scale bar: 50umThe bottom set of panels show magnified axonal segments with GFP-GSDME, mKate-OMP25 and merged channels. Scale bar: 10um (B) Quantification of the number of GFP-GSDME puncta and percentage of mitochondria enriched in GFP-GSDME, from neurons 48h after transfection with either iRFP (control), PR-50 or TDP-43 plasmids. (C) Immunoblots of primary mouse neurons transduced with lentiviruses encoding PR-50, GFP, TDP-43 or no virus. Lysates were collected at 72h post-transduction. (D-E) Primary mouse neurons were stained with TMRM and transduced with lentivirus encoding either (D) PR-50 or (E) TDP-43. 20X images were taken every 4h, starting 24h post-transduction. TMRM intensity per well was quantified and normalized relative to baseline (24h post-transduction) levels. Data points represent an average ± SEM of two independent experiments (n=2) with each condition containing four technical replicates per condition. (F-G) Representative Tuj1 staining from of wild-type and GSDME knockout primary mouse transduced with lentivirus encoding either GFP control, (G) PR-50 or (H) TDP-43. Neurons were fixed and staind for Tuj1 four days post-transduction. (H-I) Quantification of microtubule depolymerization index from mouse neurons transduced with GFP control, (H) PR-50 or (I) TDP-43 and stained for Tuj1. Violin plots display the combined median and interquartile ranges for depolymerization index taken from three independent experiments. (J-K) WT and GSDME KO mouse cortical neurons were transduced with lentiviruses encoding (J) TDP-43 or (K) PR-50 and assessed for LDH release at 4d post-transduction. For datasets D-E, two-way ANOVA (row factor = time, column factor = genotype) was performed, followed by multiple comparisons for each timepoint comparing WT vs GSDME KO groups (adj p-values, Tukey method). For datasets H-I, two-way ANOVA (row factor = toxin dose, column factor = genotype) was performed, followed by multiple comparisons for each group. p-values are indicated above brackets comparing respective datasets.

### TDP43 and PR-50 induced mitochondrial damage and neurite loss are GSDME-dependent

We next asked whether GSDME could contribute to neuronal dysfunction caused by ALS/FTD associated proteins in mouse neurons. We found that WT neurons transduced with TDP-43 and PR-50 exhibited both significant neurite loss and mitochondrial depolarization indicated by loss of TMRM staining at 72h and 96h post-treatment (**Fig. 6D-E** and **Fig S12B**). GSDME KO cells had reduced mitochondrial depolarization compared to WT neurons (**Fig 6D-E** and **Fig S12B**). These experiments indicate that ALS/FTD associated proteins can cause GSDME activation, mitochondrial localization and mitochondrial depolarization that partially relies on GSDME.

Human iNeurons are often used to model disease *in vitro* ^35^, and express full-length GSMDE at baseline (**Fig S8A**). To ask whether neurodegeneration-associated proteins could activate GSDME in human cells, we co-transfected cortical iNeurons with plasmids encoding WT TDP-43 (or iRFP control), GFP-GSDME and mKate-OMP25. TDP-43 transfected neurons exhibited robust GFP-GSDME puncta that colocalized with mitochondria (**Fig S12C-D**). As in mouse neurons, we found that lentiviral transduction with FLAG-tagged TDP-43 (at MOI 4 and 8) led to cleavage of endogenous GSDME relative to GFP transduced controls (**Fig S12E**). When taken together, these data suggest that a diverse range of neurotoxic stimuli—mitochondrial toxins, PR-50 and TDP-43 can lead to downstream GSDME activation and mitochondrial colocalization in both mouse and human neurons.

Given that GSDME KO cortical neurons had partially rescued mitochondrial depolarization, we assessed whether these neurons would also have spared neuritic processes. We took two complimentary approaches to answer this question. First, we transfected neurons with GFP and either TDP-43 or PR-50 plasmids. We then measured neurite area of transfected neurons using the GFP expression as a marker. Compared to WT neurons, GSDME KO cells transfected with PR-50 or TDP-43 had preserved neurite area (**Fig S12F-I**). Next, we used lentiviruses encoding either PR-50 or TDP-43 and measured resulting microtubule depolymerization index in cortical neurons at 72h (TDP-43) or 96h (PR-50) post-transduction. GSDME KO neurons were protected from neurite loss, with results mirroring those obtained using mitochondrial toxins (**Fig 6F-I**). At these timepoints there was minimal LDH release from WT and KO cells compared to untransduced controls (**Fig 6J-K**). This indicates that TDP-43 and PR-50 mediate GSDME-dependent axonal loss occurs prior to frank cellular necrosis. Collectively, these experiments suggest that ALS/FTD proteins engage GSDME to drive neuritic mitochondrial dysfunction and axon loss.

### GSDME impacts disease progression in the SOD1^G93A^ ALS mouse model

We next tested the hypothesis that GSDME could drive neuronal dysfunction in an animal model of neurodegeneration. ALS has been linked to mutations in the human superoxide dismutase 1 (SOD1) gene. Human SOD1^G93A^ transgenic mice develop progressive motor neuron loss in the ventral spinal cord and paralysis^49^. Mutant SOD1 proteins have been shown to induce mitochondrial dysfunction, impaired ATP production, inefficient calcium buffering and increased apoptosis of motor neurons^28, 29, 50^. To assess the relevance of GSDME *in vivo*, we first probed spinal cord lysates from the SOD1^G93A^ mouse model of ALS for GSDME activation. We detected GSDME cleavage products in spinal cords of symptomatic SOD1^G93A^ mice (P140), which was further increased by late-stage (P160) (**Fig 7A-B, Fig S13A-B**). Cleaved GSDME was not detected pre-symptomatic P82 animals (**Fig S13A**). Given this activation pattern, we hypothesized that GSDME may contribute to disease progression in the SOD1^G93A^ model. We created mice heterozygous for the SOD1^G93A^ transgene and either wild-type (SOD1^G93A^ GSDME WT) or knockout for GSDME (SOD1^G93A^ GSDME KO). We confirmed that experimental animals had similar copy numbers of the SOD1^G93A^ transgene irrespective of GSDME status by RT-qPCR; non-transgenic animals did not express SOD1^G93A^ (**Fig S13C**).

**Figure 7:**
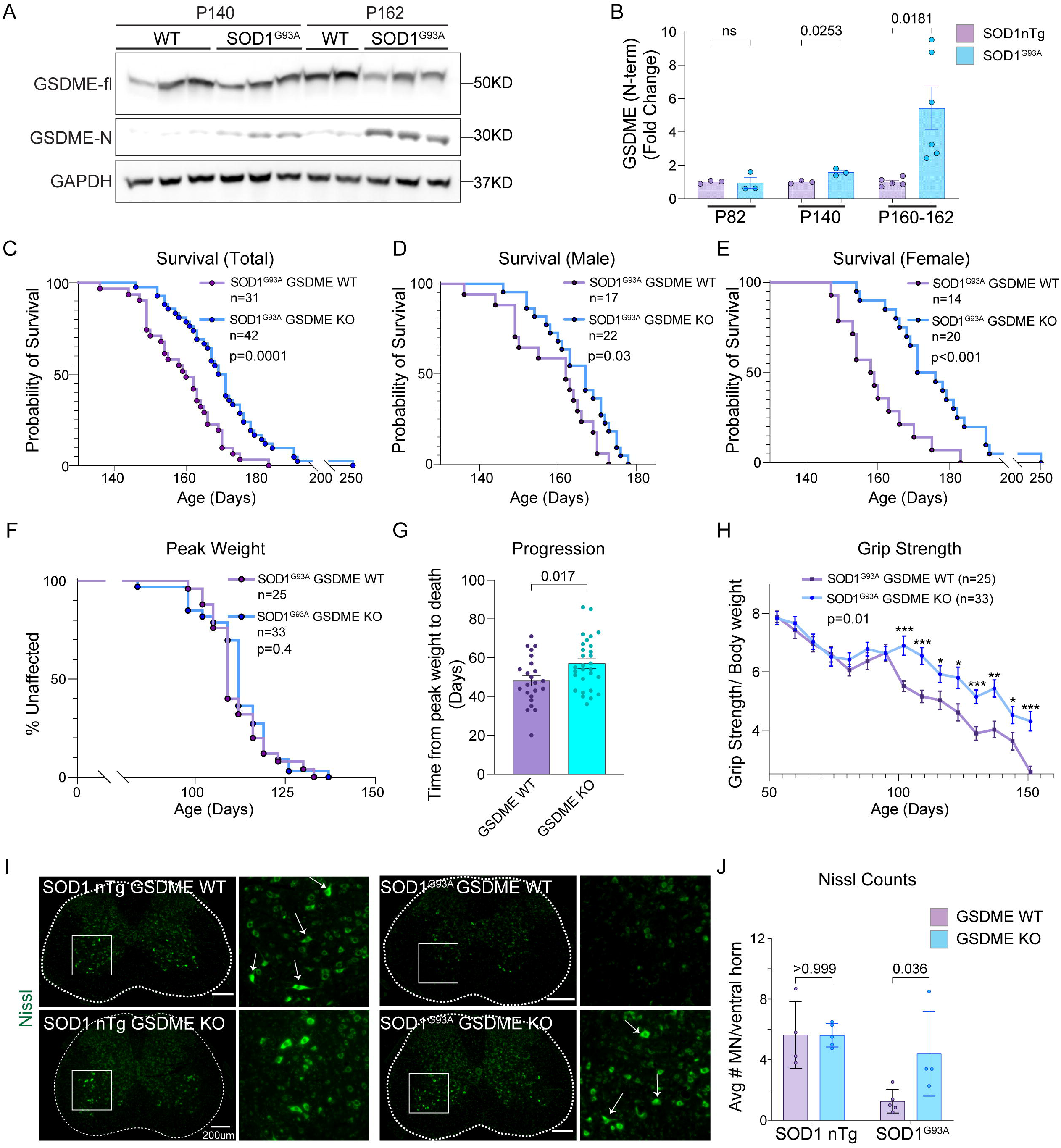
GSDME knockout rescues SOD1^G93A^ pathology in a mouse model of ALS. (A-B) Immunoblots of spinal cord lysates from SOD1^G93A^ transgenic mice and WT controls were harvested at (A) end-stage (P162) and symptom onset (P140). (B) The levels of GSDME N-terminal (normalized to GAPDH) was quantified (n=3-6) for each timepoint and genotype. (C-E) Transgenic (C) mixed gender mice that were either SOD1^G93A^ GSDME WT (N=31 total) or SOD1^G93A^ GSDME KO (N=42 total) were followed for survival. These Kaplan-Meier curves are displayed separately for (D) male SOD1^G93A^ GSDME WT (N=17) and SOD1^G93A^ GSDME KO (N=22) and (E) female SOD1^G93A^ GSDME WT (N=14) and SOD1^G93A^ GSDME KO (N=20) animals. Mice were monitored daily following onset of motor symptoms ∼d140. (F-G) The (F) age at which maximum weight was achieved was recorded for each mouse, and used to construct a Kaplan-Meier plot. (G) The time from maximum weight until euthanasia/death was also recorded as a measure of disease progression. (H) To assess motor function, grip strength values over time were tracked for each mouse and normalized by bodyweight. (I-J) Representative images of lumbar spinal cord sections from SOD1^G93A^ GSDME WT and SOD1^G93A^ GSDME KO animals at P150 (I) were stained with fluorescent Nissl dye to visualize motor neurons. (J) Quantification of the number of Nissl+ motor neurons per ventral horn of the spinal cord was performed. Each dot represents the average of 6-8 lumbar spinal cord sections from a single mouse. Magnified regions (white box) delineate a ventral horn area of a spinal cord section. For datasets C-F, Kaplan-Meier curves were constructed, and survival analysis was performed. Groups were compared for statistical differences using the log-rank test (Mantel-Cox). For datasets G and J, student’s t-tests were performed for comparisons between genotypes, and p-values were adjusted using the Tukey method. For dataset H, a two-way ANOVA (row factor = time, column factor = genotype) was performed (p = 0.01), followed by multiple comparisons for each timepoint (p-values adjusted by the Tukey method). p <0.001 = ***, p <0.01 = **, p <0.05 = ***

We found that knockout of GSDME significantly extended survival and delayed disease progression in mice containing the SOD1^G93A^ transgene (**Fig 7C**). Mean survival for SOD1^G93A^ GSDME WT mice was 159±1.9 days, while that of SOD1^G93A^ GSDME KO counterparts was 171±2.5 days (**Fig 7C**). Female SOD1^G93A^ GSDME KO animals received a slightly greater survival benefit relative to female SOD1^G93A^ GSDME WT mice compared to males (**Fig 7D-E & Fig S13F**). As a measure of disease on set, the age to maximum body weight were unaffected by GSDME genotype in SOD1^G93A^ animals (**Fig 7F & Fig S13D**). However, disease progression—as measured as time from maximum body weight until the time point of euthanasia—was significantly extended in SOD1^G93A^ GSDME KO animals relative to SOD1^G93A^ GSDME WT littermates (**Fig 7G**). Loss of grip strength was also significantly delayed in SOD1^G93A^ GSDME KO compared to SOD1^G93A^ GSDME WT mice (**Fig 7H**). No baseline differences in grip strength were recorded for non-SOD1 transgenic GSDME WT and GSDME KO animals (**Fig S13E**). These results indicate that GSDME deficiency impacted the symptomatic progression phase in this animal model, leading to an overall motor and survival benefit (**supplementary video 4**).

Consistent with the rescue in grip strength and survival, motor neurons in the ventral horn were preserved in SOD1^G93A^ GSDME KO mice at P150 (late-stage) relative to age-matched SOD1^G93A^ GSDME WT littermates (**Fig 7I-J**). We also looked at makers for glial activation in the spinal cord, as microgliosis and astrogloisis are hallmarks of disease progression in the SOD1 mouse model of ALS and in human ALS patients^51–55^. We observed that GSDME KO mice showed decreased intensity of GFAP+ astrocytes (astrogliosis) in the ventral horns of SOD1^G93A^ animals (**Fig S14A-B**). SOD1^G93A^ GSDME KO animals also showed decreased microgliosis, as measured by the number of Iba1+ positive cells in the ventral horn (**Fig S14C-D**) and levels of the microglial lysosomal activation marker, CD68, compared to those in SOD1^G93A^ GSDME WT mice (**Fig S14E-F**). When taken together, this data demonstrates that GSDME is activated in a classic mouse model of ALS and drives pathology—namely neuronal loss and gliosis—that contributes to decreased motor function and survival.

### GSDME mediates neurite loss in ALS patient IPSC-derived motor neurons

To determine potential relevance to human disease, we mined genome-wide expression profiling data of laser capture microdissection-enriched motor neurons (MNs) from patients with sporadic ALS (sALS)^56^. We found that GSDME was significantly upregulated in neurons from sALS patients relative to age-matched healthy controls (**Fig S15A**). This data indicates that GSDME protein is present in the human CNS and may be relevant in the setting of neurologic disease.

We next investigated whether GSDME may contribute to neuronal dysfunction in human ALS patient iPSC-derived motor neurons. To model human neurodegeneration *in vitro,* we cultured a control iPSC-derived MN line (1016A) and an ALS iPSC-derived MN line (*TDP43*^G298S^). Challenge of these iMNs with a proteasome inhibitor (MG132) or ER-stress toxins (thapsigargin and tunicamycin) led to dose dependent neurite loss and cell death; these effects are exacerbated in the mutant *TDP43*^G298S^ line (**Fig S15-B-E**)^57^. We next determined if knockdown of GSDME expression in human iPSC-derived motor neurons would rescue this ER-stress driven neurite loss We next found that *TDP43*^G298S^ motor neurons transduced with lentivirus encoding scrambled (control) shRNA were highly vulnerable to neurite destruction caused by MG132, tunicamycin or thapsiagargin (**Fig 8A, Fig. S15B-E**). By contrast, knockdown of GSDME using lentiviral delivery of two targeting shRNAs (sequence #1 and #3) led to significant sparing of neurite loss in *TDP43*^G298S^ iMNs (**Fig 8, S15 F-G**). Rescue due to GSDME knockdown was also observed in control 1016A (WT) iMNs treated with ER toxins or MG132—though to a smaller extent than in susceptible *TDP43*^G298S^ iMNs (Fig S15 A-E). Collectively, these results support that GSDME drives pathology downstream of ER-stress and proteasome inhibition in WT MNs, and to a much greater degree, in the vulnerable *TDP43*^G298S^ genetic background. Thus, GSDME deficiency can rescue neurite loss in a patient-derived ALS model.

**Figure 8:**
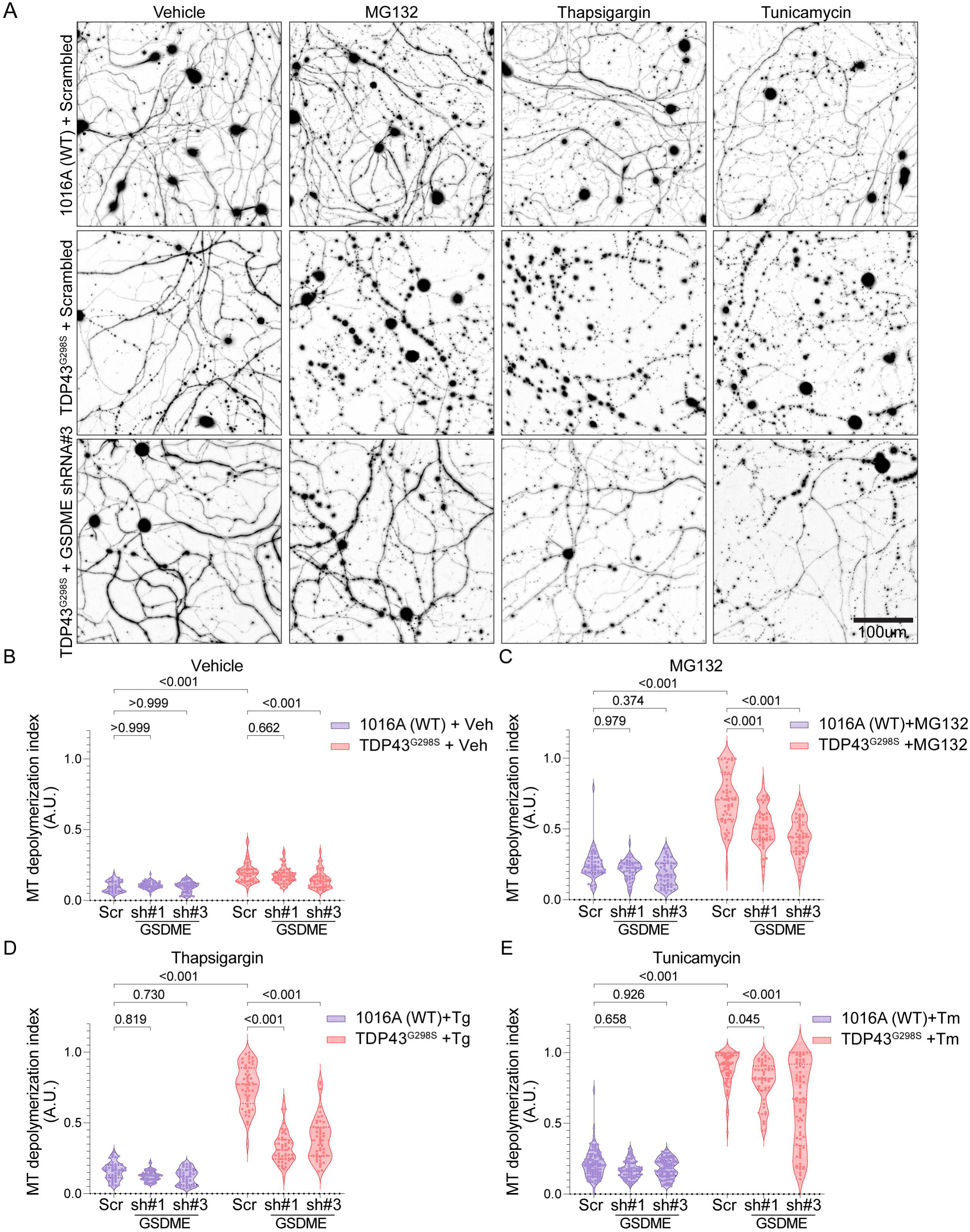
GSDME knockdown rescues neurite loss in ALS patient iPSC-derived motor neurons. (A) Representative images of control 1016A or *TDP43*^G298S^ iPSC-derived motor neurons transduced with either scrambled shRNA or GSDME targeting shRNA #3 and treated with either DMSO, tunicamycin, MG132 or thapsiagargin and stained for Tuj1 at 48h post-toxin treatment. (B-E) Microtubule depolymerization index was calculated for 1016A (WT) and *TDP43*^G298S^ motor neurons treated with (B) 0.1% DMSO (vehicle) (C) 1uM MG132 (D) 0.5uM of thapsiagargin (E) 5uM tunicamycin and either scrambled shRNA or GSDME targeting shRNA#1 or #3. Violin plots display the combined median and interquartile ranges for depolymerization index. For datasets B-E two-way ANOVA was performed, with multiple comparisons for each group (adj p-values, Tukey method). For B-E images were collected from 6 technical replicates (wells) ± SEM.

## Discussion

Mitochondrial dysfunction is an important and early hallmark of neuronal injury. Here we describe GSDME as a critical regulator of mitochondrial damage, neurite loss, and death in neurons downstream of multiple classes of neurotoxic stimuli. Both mouse and human neurons express GSDME, and this protein is activated following exposure to mitochondrial toxins and ER stressors. Several such toxins (e.g. rotenone and MPTP) are linked to neurodegeneration in humans, and others can be used to model features of disease *in vitro* and in mouse models ^58, 59^. GSDME is also activated following neuronal expression of ALS/FTD associated proteins, TDP-43 and PR-50. Activated GSDME rapidly colocalizes with axonal mitochondria prior to overt cell death, driving both mitochondrial damage and neurite loss. We further show that GSDME plays a significant role in the SOD1^G93A^ mouse model of ALS and in an iPSC derived motor neuron model of neurodegeneration, thus identifying this axis as a therapeutic target.

We uncover several aspects of GSDME biology unique to neurons that differ from previous studies in cell lines or primary immune cells. We show that GSDME mediates axonal mitochondrial damage and local neurite loss. In neurons, active GSDME primarily localizes to mitochondrial membranes and mediates mitochondrial destruction and neurite loss, in the absence of significant plasma membrane localization and rupture (as assessed by LDH release). There may be multiple reasons behind the resistance of neurons to membrane rupture. For example, the neuronal plasma membrane lipid composition may be uniquely resistant to N-GSDME insertion, or may harbor components that limit its activity in neuronal cell membranes (akin to the relationship between XIAP and CASP3). It is also possible that neuronal plasma membrane damage from GSDME activation is rapidly repaired, while delicate axonal processes are not spared. Such a phenomenon – ESCRT complex repair of gasdermin-mediated plasma membrane damage - has been described following GSDMD activation in myeloid cells^60, 61^. In macrophages, this is important for cytokine secretion as it allows “hyperactivated” living macrophages to engage GSDMD mediated IL-1 release without overt cell death ^60, 61^. In the CNS, there may be a physiologic benefit in destroying distal processes while keeping the soma intact. For example, GSDME mediated neurite pruning may be crucial to either development or host defense (e.g. to stop the spread of viral or microbial pathogens). However, in the setting of progressive neurodegenerative pathology (such as in ALS), GSDME may be inappropriately activated to drive disease progression and contribute to neuroinflammation. Our findings thus expand the biological role for GSDMS in the CNS. Elucidating the physiological contexts (e.g. development or infection) of GSDME mediated neurite loss would be an important future direction.

Prior work explored GSDME activation in cancer cell lines, focusing primarily on its contribution to overt cellular necrosis^11, 25, 38^. Activated caspase-3 cleaves and activates GSDME, converting noninflammatory apoptotic cell death to inflammatory pyroptosis. GSDME colocalizes with mitochondria in HeLa cells treated with DNA-damaging agents (e.g. UV radiation, etoposide) and mediates cytochrome-c release to further amplify caspase-3 processing and necrosis^10, 38^. Oxygen or glucose deprivation also leads to caspase-1/Bid/caspase-3 activation in cortical neurons; it has been suggested that GSDME may contribute to cytochrome-c release and cell death in this context^62^. More recent work in a mouse model of macular degeneration and a murine retinoblastoma cell line has implicated GSDME in amplifying cytochrome-c release and capsase-3 activation in sensory neuroepithelium^23^. We establish this positive feedback loop in primary neurons and map its neuron-specific cell biology—namely localized GSDME activation in axons, mitochondrial damage, disruption of mitochondrial trafficking and neurite-loss.

Spatially restricted GSDME activity in axons (**Figs 3 & 5**) raises the possibility that GSDME is a key downstream effector of caspase-3 mediated axonopathy and neuronal demise. Caspase-3 inhibition has been shown to protect neurons from age or disease related axon loss and cell death^3, 63–65^. In neurodegeneration, a unique combination of environmental exposures and gene expression differences may predispose certain neurons to activate the caspase-3/GSDME axis. How different neuronal cell types reach this terminal point—dopaminergic neurons in PD, motor neurons in ALS and cortical neurons in FTD—remains an active area of investigation.

Neurons may utilize multiple pore-forming molecules to regulate mitochondrial membrane integrity; namely upstream pro-apoptotic BCL-2 family members (e.g. Bax, Bak, Bid), the mitochondrial permeability transition pore (MPTP) and in concert with GSDME. Caspase-3 cleaves GSDME, allowing for N-terminal oligomerization and insertion into lipid membranes. Following mitochondrial colocalization, GSDME subsequently drives mitochondrial depolarization, cyt-c release, and enhanced caspase-3 activation. Thus, GSDME is a versatile lipid-targeting pore forming molecule that may have overlapping functions with Bax/Bak families as well as MLKL—the pore-forming executioner of necroptosis ^66, 67^. How these distinct families of pore-forming molecules operate in a cell-type specific manner in neurodegeneration, or work in concert to form compensatory signaling loops is poorly understood, and an important area of future investigation. Given the link between mitochondrial toxins and stress to quality control programs (mitophagy and mitochondria derived vesicles), it would also be important to see how these mechanisms function either in series or in parallel to GSDME activation.

We found that expression of the active N-terminal GSDME fragment in neurons is sufficient to cause mitochondrial dysfunction and neuritic retraction independent of any other stimuli, indicating that it may have a preference for mitochondrial membranes in neurons. Indeed, biophysical assays have shown that N-terminal GSDME (as well as GSDMD) binds cardiolipin with much higher affinity than it does phosphatidylcholine, phosphatidylinositol or other lipids on the OMM ^11, 13^. Mitochondrial stress can externalize cardiolipin from the inner to outer mitochondrial membrane^68, 69^. Our work demonstrates that in neurons, GSDME rapidly localizes to mitochondria prior to axon degeneration and neuronal cell death. Knockdown of cardiolipin synthase (CLS1) reduces GSDME colocalization with mitochondria, suggesting a role for cardiolipin in homing of GSDME to mitochondria. However, this mainly occurred at early time points (**Fig S7**)—it is unclear whether this partial rescue is due to incomplete knockdown or the potential ability of GSDME to bind other ligands present in the OMM. Investigating which other lipids or proteins regulate GSDME binding to the OMM will be an important avenue of future mechanistic investigation.

Destruction of axons is an early step in conditions such as neurodegeneration, neuropathic pain, and stroke ^26, 27, 70^. Our findings using imaging and microfluidic compartments demonstrate that GSDME activation can occur locally in axons and drive local mitochondrial damage, without involving the cell body. Thus, our study positions GSDME as a molecule that may work alongside calpains, SARM1, KIF2A, MAPK-JNK, loss of STMN2 and other effectors to disrupt axons. Though phosphorylation and palmitoylation of GSDME residues can modulate its activity, there are no known GSDME regulators apart from upstream proteases (caspase-3 and granzyme-B) ^25, 38, 71^. Calpains are proteases with cleavage activity linked to axon degeneration^72^. SARM1 has been shown to regulate the extent of GSDMD-driven pyroptosis in immune cells ^73^. The MAPK-JNK signaling axis promotes SARM1 dependent axonal degeneration *in vivo,* but is also an upstream regulator of GSDME activity in cancer cell lines^74–76^. It will be important to determine if SARM1, calpains, JNK or other mediators of axon degeneration regulate GSDME function in a common pathway to destroy neuronal processes.

We further showed that GSDME is activated by neurodegeneration associated proteins (TDP-43 and PR-50). That both mitochondrial toxins and aggregated proteins evoke GSDME signaling is perhaps not unexpected. TDP-43 has been shown to bind and damage mitochondria, leading to release of DNA and activation of STING signaling to drive cell death^45^. *C9ORF72* DPRs directly inhibit mitochondrial complex V, leading to membrane depolarization and neuronal death ^77^. Motor neurons of patients and mice carrying SOD1 mutations, have impaired electron transport chain function, disrupted mitochondrial ATP production, inefficient calcium buffering, and enhanced caspase-3 activation^28, 50, 55^. That mutant SOD1 aggregates colocalize with mitochondria further suggests a directly toxic effect^29^.

How neuronal GSDME gene expression is regulated remains to be determined. Transcriptional data from micro dissected motor neurons showed an upregulation of GSDME in patients with ALS compared to age-matched controls. Prior work has shown that tumor suppressor gene p53 can bind to intron 1 of the GSDME gene to induce its expression, and p53 knockout mice display reduced levels of GSDME in the GI epithelium ^19^. A recent study showed that p53 drives *C9ORF72*-dependent cortical neuron death in a mouse model of FTD/ALS ^39^. As suggested for other tumor suppressor genes (e.g. ATM, p53, Bax and WWOX), GSDME may in one context suppress cancer growth and in another, inappropriately promote neuronal injury and inflammation^78–80^.

When we knocked out GSDME in the SOD1^G93A^ transgenic background, we observed a slowing of disease progression. This data functionally links the activation of GSDME with neurodegeneration in a mouse model of ALS. We found GSDME was activated (cleaved) in spinal cords at late stages of disease and its deficiency led to decreased microglia and astrocyte activation. Previous studies have found that activation of GSDME impacts mouse models of tumor progression, chemotherapy induced lung damage, cisplatin-induced kidney injury, macular degeneration and inflammatory bowel disease ^22, 23, 25, 81, 82^. In these settings, knockout of GSDME not only reduces cell death, but also reduces local inflammation. Mapping how exactly GSDME promotes *in vivo* neuroinflammation—either via neuronal release of pro-inflammatory cytokines, damage-associated molecules (e.g. HMGB1 and ATP), or other secreted factors to recruit glia or CNS-infiltrating immune cells—will be an important step in understanding its role in disease pathophysiology.

We extended our *in vivo* results by employing an *in vitro* human model of ALS—assessing neurite loss in ALS iPSC-derived motor neurons that carry a TDP-43 mutation (**Fig 8**). In these cells, neurite loss was rescued by GSDME knockdown, supporting that this pore-forming molecule may contribute to ALS pathology. These iPSC-derived *in vitro* systems are important tools to further understand the mechanisms by which GSDME is activated in human disease and affects neurodegeneration.

In conclusion, our work positions GSDME as an intrinsic executioner of axon degeneration and cell death in neurons. When activated by diverse neurotoxic stimuli, GSDME drives features of neurodegeneration *in vitro*—destruction of mitochondria, enhancement of caspase-3 activation and neuritic retraction—and contributes to disease progression in two ALS models. Recently, small-molecule inhibitors of GSDMD activation have been identified ^83, 84^, though it is unclear whether GSDME is similarly druggable. Future work should determine the feasibility of targeting GSDME expression or function. Ultimately, determining how multiple cell death axes— gasdermins, caspases, calpains, and necroptotic machinery—contribute to neuronal cell death may suggest new therapeutic strategies for neurodegenerative diseases.

### Limitations of this study

*In vitro* analysis using mouse and iPSC-neurons allows for well controlled pharmacological and genetic manipulation, careful microscopic examination and discovery of exciting new aspects of neuronal cell biology that are not possible to observe currently *in vivo*. However, cultured mouse and human iPSC-derived neurons do not recapitulate the complexity of the nervous system. *In vivo*, it is possible that there are intrinsic compensatory feedback loops and non-cell autonomous mechanisms influence GSDME signaling. Such interactions may not be possible to model in pure neuronal cultures. Further, biochemical purification of mitochondria in cultured systems may be contaminated by plasma membrane fractions. Additional experiments controlling for this possibility, such as neuronal plasma membrane fractionation, would strengthen the work. We demonstrate that GSDME is engaged in neurons after toxin or FTD-associated protein exposure, in several cellular contexts lending strength to our conclusions. Future studies employing mouse models of ALS/FTD coupled with GSDME knockout will be necessary to further elucidate the role of GSDME in a more physiologic context of neurodegeneration. Lastly, experiments showing *in vivo* human GSDME cleavage and immunostaining are inherently limited by time resolution—cell death proteins are difficult to detect *in vivo* due to their transitory nature. Once a cell dies, its mRNA and protein may not be readily detectable by standard methodologies. Increased sampling of tissues from mouse models and human patients will elucidate when and how this molecule is regulated during disease.

## Acknowledgements

We would like to thank Dr. Aaron Gitler for providing plasmids encoding PR-50 and TPD-43, and Dr. Laura Volpicelli-Daley and Dr. Ulf Dettmer for advice and collaboration on related work. We would also like to thank Dr. Nandini Ramesh for helping to secure mouse brain samples. We would like to acknowledge Dr. Thomas Schwarz, Dr. Beth Stevens and Dr. Francisco Quintana for their feedback and helpful discussions regarding preparation of this manuscript. Lastly, we would like to thank Alex Yueting-Lu, Victoria Tong and Samantha Choi for excellent technical support during this project. This work was supported by R01DK127257 (IC); Chan-Zuckerberg Institute Neurodegeneration Challenge Network (IC); R01CA240955 (JL); Cancer Research Institute fellowship (RM); R01AG055909 (TYP); R01NS115139 (ACP); U19 AG062418 and P50 NS053488 (UPenn Brain Bank, EL). ACP is additionally supported by the Parker Family Chair, the Chan Zuckerberg Initiative Neurodegeneration Challenge Network, and the AHA/Allen Brain Health Initiative. XJ is supported by a Development Grant from the Muscular Dystrophy Association. CLT is the recipient of the Araminta Broch-Healey Endowed Chair in ALS. The project described was also supported by award numbers T32GM007753 (NIGMS) and F31NS122292 (NINDS) (DVN).

## Author Contributions

Conceptualization: DVN, HB, IC

Experimental data collection and analysis: DVN, HB, HC, GG, MB, RM, EB and ACP, VC

Securing key reagents: LP, YJZ, TYP, CLT, JL,

Writing original draft: DN, HB, IC

Review and Editing manuscript: DN, HB, IC, GG, ACP, JL, TYP, CLT

Funding acquisition: IC

## STAR Methods

### Animals

All mouse experimental procedures were performed in compliance with the Harvard Medical School and The Jackson Laboratory Institutional Animal Care and Use Committees. C57BL/6NJ (# 005304), C57BL/6N-*Gsdme^em1Fsha^*/J (# 032411), and SOD1^G93A^ mice (JAX #002299) mice were obtained from Jackson labs and bred at Harvard Medical School. Animal experiments were fully approved by the Harvard Medical School Institutional Animal Care and Use Committee (IACUC). Animals were housed in temperature (22 ± 2 °C) and humidity (55 ± 5%) controlled care facilities at Harvard Medical School on a 12 h light:dark cycle,and provided with freely available food and water.

### SOD1^G93A^ Breeding and Behavioral Assessments

#### Breeding

SOD1^G93A^ mice (JAX #002299) were originally acquired from Jackson Laboratory (Bar Harbor, ME). The SOD1^G93A^ GSDME KO mice were obtained by crossing male mice carrying the SOD1^G93A^ transgene with female GSDME KO (JAX #032411) mice for 3 generations. The control SOD1^G93A^ GSDME WT mice, were obtained by crossing SOD1^G93A^ mice with wild type female mice having a B6NJ background (JAX #005304), also for 3 generations. This breeding strategy controlled for any strain background related phenotypic differences.

All offspring were tested for equivalent SOD1 gene copy numbers by qPCR (forward primer: CAGTAACTGAGAGTTTACCCTTTGGT; and reverse primer: CACACTAATGCTCTGGGAAGAAAGA. Mice that had SOD1 signal drop-off of more than 30% from control SOD1^G93A^ mice were considered as low copy number animals and were excluded from the study. Male and female mice were used for all experiments and experimental groups were sex balanced. Animals were provided with food and water *ad libitum*.

#### Weight Measurement

Weights of all mice carrying the SOD1^G93A^ transgene were measured biweekly from week 7 to week 21. Disease onset was defined as the day of peak weight while disease progression was defined as the number of days between disease onset to euthanisia.

#### Grip strength

Grip strengths of the SOD1^G93A^ and SOD1^G93A^ mice were measured weekly, starting at week 7 and ending at week 21. A grip strength machine (BIOSEB# bio-GS3) with a mesh grid attachment was used to obtain combined grip strengths of all four limbs. For each measurement, the mouse was held at the base of the tail and placed on the mesh grid. A total of 5 pulls were performed per mouse with approximately 5-10 second breaks between each pull. The top three pulls per time point were averaged and used for data analysis. 12 or more mice per genotype, per sex, were used for weights and grip strength analysis.

#### Survival

Each mouse was monitored daily after symptom onset. Mice that were unable to rear were given hydrogels and wetted food pellets at the bottom of the cage. Euthanasia time points for each mouse were determined by the inability to successfully right itself when flipped on either of its sides within 30s.

### Brain and spinal cord tissue lysis

For analysis of mouse CNS tissue, animals were anesthetized with Avertin solution (500 mg kg−1, MilliporeSigma) and perfused with 20 ml of cold PBS before harvest. Brains and spinal cords were dissected in dish of cold PBS and stored at −80 C prior to tissue homogenization/lysis. Lysis buffer was prepared on ice as follows: 10 ml T-per buffer (Thermo #78510), 1 tablet protease inhibitor (Sigma #11836153001/Roche), 100ul HALT protease inhibitor (Thermo #87786), 100ul 0.5M EDTA (Thermo #87786), 1 tablet Phosstop™-phosphatase inhibitor tablets (Sigma #04906845001/ Roche). Tissue lysis was performed as previously described ^85^. Briefly, dissected and frozen tissue was resuspended in cold lysis buffer in a 2mL Eppendorf tube. One autoclaved metal bead (BB) was placed in each tube containing sample, and lysed in a bead beater (Qiagen TissueLyser II) at a setting of 25 Hz for 5 minutes. Homogenized tissue samples were then incubated on a rotating tube rack at 4C for 30 min. Following incubation samples were clarified by spinning at 16000 x g for 15mins at 4C. Supernatants (lysate) were then taken and stored at - 80 C prior to analysis. For immunoblot processing, sample lysates were diluted in Bolt 4X loading buffer (Thermo #B0007) containing 1X Bolt Sample Reducing Agent (Thermo #B0009) and beta-mercaptoethanol. Samples were then boiled for 10 min at 90 C, spun down and stored at −20 C.

### Primary mouse neuron culture and plating

Mouse (C57BL/6NJ or C57BL/6N-*Gsdme^em1Fsha^*/J) primary cortical neurons were dissociated and cultured using the Worthington Papain Dissociation System (Worthington Biochemical, cat # LK003153). Briefly, passage 0 (P0) mouse cortices were dissected and collected in cold Earle’s Balanced Salts (EBSS). Cortices were then resuspended in 2.5 ml of warmed EBSS with papain (20 units ml^-1^) and DNase (2000 units ml^-1^). Following a 12 min incubation at 37°C, cortices were triturated 12 times using a 10mL glass pipette, and then 12 times using an 18G needle and syringe to make a single cell suspension. Samples were then passed through a 70um mesh filter to remove debris, and the filtrate was centrifuged (2000xg for 5 min) to pellet cells. Cells were then resuspended in 1.6 ml of suspension media [1.375 ml EBSS, 150 µl albumin-ovomucoid inhibitor (10 mg ml^-1^ in EBSS), and 75 µl DNase (2000 units ml^-1^)]. This solution was layered on top of a 2.5 ml solution of albumin-ovomucoid inhibitor (10 mg ml^-1^ in EBSS) to create a continuous density gradient, and the samples were centrifuged at 1000 rpm for 5 min. The supernatant was discarded, and pelleted neurons were collected in warm Neurobasal^TM^ Plus medium (Thermo Fisher Scientific) supplemented with 200 mM L-Glutamine with 1% (v/v) penicillin-streptomycin. For plating of primary neurons, dishes were pre-coated with 20 ug/mL poly-L-lysine (Sigma Cat# P2636-25MG) and 6ug/mL laminin (Gibco Cat# 23017-015) in ddH20 for 2h at 37C. Dishes were washed twice with ddH20 immediately before plating neurons. For GFP-GSDME imaging experiments, cells were plated in 24 well glass bottom dishes (cellVis: P24-1.5H-N) at a density of 3.5 x 10^5^ cells per well. For immunoblots, cells were plated at a density of 1 x 10^6^ cells per well in 6 well plates (cellVis: P6-1.5H-N). For cell death, TMRM assays and Tuj1 staining, neurons were plated in 96 well glass bottom dishes (Cellvis #P96-1.5H-N) at a concentration of 5 x 10^4^ cells/well.

### SH-SY5Y culture

Wild-type and GSDME -/- SH-SY5Y cell lines were a gift from Dr. Judy Lieberman (Boston Children’s Hospital). SH-SY5Y were cultured in DMEM medium supplemented with 10% heat-inactivated fetal bovine serum (FBS), 6 mM HEPES, 1.6 mM L-glutamine, 50 μM 2-mercaptoethonal, 100 U/ml Penicillin G and 100 μg/ml streptomycin sulphate. CRISPR-Cas9 knockout of GSDME in SH-SY5Y was performed as previously described.^25^ Briefly, *GSDME* gRNAs (5’-TAAGTTACAGCTTCTAAGTC-3’ and 5’ -TGACAAAAAAGAAGAGATTC-3’) were cloned into LentiCRISPR-v2 puro vector. The resulting plasmids were transfected into HEK293T with pSPAX2 and pCMV-VSV-G at a 1:1:2 ratio. Supernatants containing lentivirus were collected 2 days later and used to transduce SH-SY5Y at an MOI of 0.3. Two days after transduction, 3 μg/ml (8 μg/ml for CT26) puromycin or 200 μg/ml hygromycin was used to select for positive cells (5d incubation). Cells were then subcloned by limiting dilution in 96-well plates and screened for GSDME expression by immunoblot. LentiCRISPR-v2 empty vector was used to generate control cells.

### Human iPSC-derived cortical neuron culture and plating

NGN2 tetracycline-inducible iPSC neuron lines were a gift from the Tracey Young-Pearse lab and were generated using a lentiviral transduction protocol as previously described ^86^. Plates were coated with poly-orinatine and laminin (10 ug/mL poly-ornithine, 5 ug/mL laminin) 24 hours prior to cell plating. Plates were coated with Matrigel basement matrix 2 hours prior to cell plating (8.7 µg/cm2). Day 4 cells were thawed from stock and plated at 150,000 cells per well with iN media (Neuralbasal media, 1% Glutamax, 0.3% Dextrose, 0.5% MEM NEAA, 2% B27, 10 ng/mL of each BDNF, GDNF, and CNTF, 5 ug/mL puromycin, 2 ug/mL doxycycline, 10 uM Y-27632). Half media changes with the same iN media without Y-27632 were performed every 2-3 days prior to harvest.

Specific reagents used for culturing iPSC derived neurons are as follows:

| Reagent | Vendor | Catalogue Number |
| --- | --- | --- |
| Poly-ornithine | Sigma | P3655-50mg |
| Laminin | Life Technologies | 23017-015 |
| Matrigel® Basement Membrane Matrix | Corning | 354234 |
| Neurobasal Medium | Gibco | 21103 |
| Glutamax | Gibco | 35050 |
| Dextrose | Sigma | D9434-250g |
| MEM NEAA | Invitrogen | 11140-050 |
| B27 | Life Technologies | 17504-044 |
| BDNF | Peprtech | 450-02 |
| GDNF | Peprtech | 450-13 |
| CNTF | Peprtech | 450-10 |
| Puromycin | Life Technologies | A11138-03 |
| Doxycycline hyclate | Sigma | D9891-5g |
| ROCKi (Y-27632) | Stemcell Technologies | 72304 |

### Drugs and treatments

Raptinal (cat# SML1745), rotenone (cat# R8875), antimycin-A (cat# A8674) and 3-Nitropropionic acid (cat# N5636) were purchased from Sigma and resuspended in DMSO to make stock solutions. For raptinal treatments, cells were exposed to final concentrations ranging from 1-10uM for 1h, and then incubated with fresh media and assessed at later timepoints. For antimycin-A treatments, cells were exposed to final concentrations ranging from 5-20 uM for 30 min and then incubated with fresh media. For rotenone and 3-NP treatments, cells were exposed to working concentrations ranging from 10-30uM (Rotenone) and 1-3 mM (3-NP) for 2h and then incubated with fresh media. Z-VAD-FMK (R&D cat# FMK001) was resuspended in DMSO and used at a final concentration of 20 uM. For experiments using toxins and z-VAD, neurons were pretreated with 20uM z-VAD for 30 min, and then treated with a combination of zVAD (20uM) and toxins for the indicated timepoints.

### Primary neuron transfection

Primary neurons were transfected at DIV 6-7 and analyzed at DIV9-10 (2-3 days following transfection). Transfections were performed using lipofectamine 2000 reagent (Thermo Scientific cat# 11668027) as previously described ^87^. Briefly DNA and lipofectamine mixes were made in plain neurobasal media (NBM, without PSG and B-27) with 0.25-1ug of DNA and 0.75-3uL of lipofectamine reagent (1:3 DNA/lipofectamine ratio). The DNA and lipofectamine mixes were gently vortexed (30s) and incubated at RT for 15 min. Prior to addition of the transfection mix, conditioned media was removed from neuronal plates and saved. Neurons were then washed 3 times with warmed plain NBM, and DNA/lipofectamine mix (100uL/well for a 24 well plate) was added to each well and incubated for 1h at 37C. Following incubation, each transfected well was washed 3 times with fresh plain NBM and the conditioned media was added back to each well. Neurons were then incubated for 2-3 days at 37C prior to imaging and analysis.

### Plasmids

All plasmids used in this study for transfections are as follows:

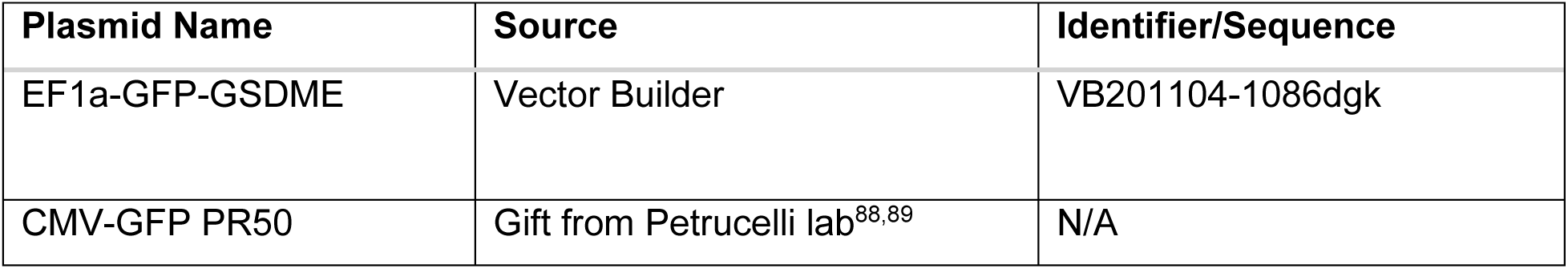
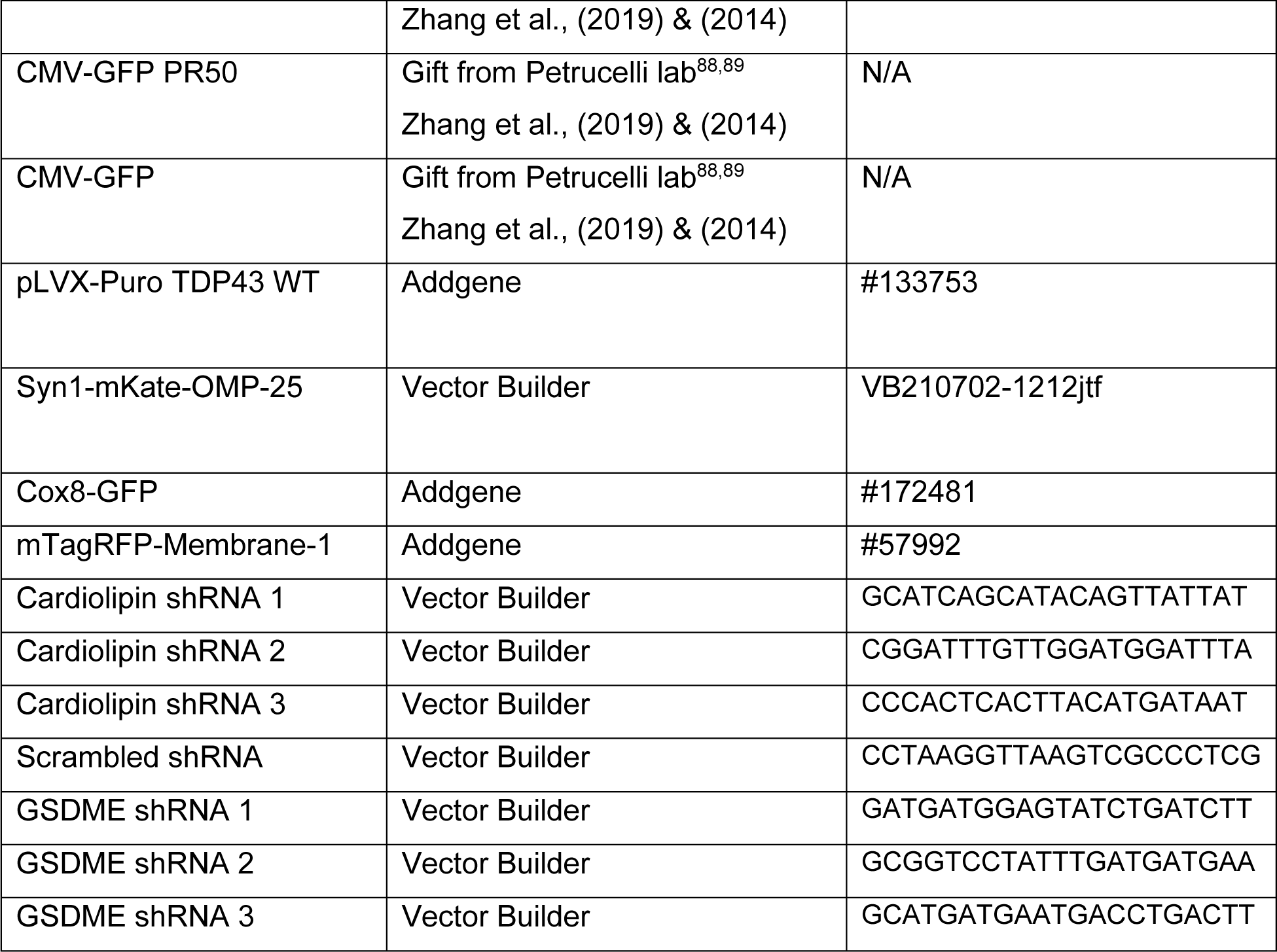

### Viruses

All viruses used in the study for transductions are as follows:

Viruses All viruses used in the study for transductions are as follows:
| Virus | Source | Identifier |
| --- | --- | --- |
| Lenti-Syn-PR50 | Gift from Gitler lab <sup>39</sup> | N/A |
| Lenti-Syn-eGFP | Gift from Gitler lab <sup>39</sup> | N/A |
| Lenti-Syn-TDP43 | Gift from Gitler lab <sup>39</sup> | N/A |

### Lentivirus transductions of primary neurons

Neurons were transduced overnight with lentiviruses encoding control (eGFP) or PR-50 and TDP-43 proteins as described previously.^39^ Briefly, primary neurons plated in 96-well plates at a density of 50-75,000 cells/well were grown for 72 hours. The neurons were then treated with tittered viruses at MOIs of 2 or 3. Transduction volume was 40uL/well. Following overnight incubation with virus, neurons were washed 3x with warmed complete NBM and followed for TMRM staining, immunoblot analysis or Tuj1+ staining as described below.

### Live cell imaging of neurons

For live-cell imaging of cortical neurons, neuronal cultures grown on glass bottom dishes at a density of 175,000 cells/cm^2^ were imaged on a DMI8 Zeiss microscope. The microscope was equipped with an environmental chamber that was supplied with humidified 5% CO2 and maintained at 37C. Images were acquired with an Andor Zyla sCMOS camera via a 20× Plan Apo objective. To monitor GSDME enrichment on mitochondria, images from neurons co-transfected GFP-GSDME and mKate2-OMP25 (referred to as mito-RFP) were captured every 15 minutes following treatment with toxins (raptinal and rotenone). To monitor mitochondrial motility, each well of neurons (350,000 cells/well) were sparsely transfected with mKate2-OMP25 (referred to as mito-RFP) and cytosolic GFP. Two days following transfection, the neurons were treated and imaged. Images of whole neurons (all visible neurites) were captured every 6s. Kymographs were generated from 3- to 5-min time-lapse movies and analyzed with a custom-written ImageJ macro for percent motility mitochondrial density, and length. High resolution 3-5 min timelapse movies were taken at 30 min, 1h and 2h post-raptinal treatment.

### Immunoblotting and antibodies

Cells were lysed on ice in 1x RIPA buffer (EMD Milipore cat# 20-188) supplemented with 0.5mM EDTA, 1X Halt Protease inhibitor cocktail (Thermo Scientific cat# 87786) and 1 Complete Mini protease inhibitor tablet (Sigma cat# 11836153001). Following a 15 min incubation in lysis buffer, cells were centrifuged at 18,000 x g for 15 min at 4C. Pellets were discarded and supernatants were diluted with 4X Laemmli buffer (to 1X final concentration) supplemented with 10X Bolt sample reducing agent (cat# B0009). Samples were then incubated on 90C heat block for 10 minutes, and run on Bolt 4-12% Bis-Tris-Plus gels (Thermo Scientific cat# NW04125BOX). Gels were transferred to nitrocellulose iBlot 2 membranes (Fisher Scientific cat# IB23001), blocked with 5% Pierce Clear Milk Blocking Buffer (Thermo Scientific cat# 37587) for 30 minutes, washed 3x with TBST (TBS, 0.05% Tween-20), and incubated overnight in blocking buffer containing primary antibody at 4C. Following antibody incubation, blots were washed 3x with TBST for 10 min, incubated with secondary antibody for 1h at RT, followed by three additional 10 min washes with TBST. GSDME and caspase-3 immunoblots were developed with Supersignal West Pico Chemiluminescent Substrate (Thermo Scientific cat# 34080) on a ChemiDoc MP or Azure 300 chemiluminescent imaging system. GAPDH immunoblots were incubated with IR-fluorophore conjugated secondary antibodies (LI-COR Biosciences cat# 926-32213) and developed and imaged on a Li-COR imaging system.

### Details of antibodies used for immunoblotting are as follows

**Details of antibodies used for immunoblotting are as follows:**
| Antibody Target | Identifier | Usage Dilution |
| --- | --- | --- |
| GSDME | Abcam #ab215191 | 1:1000 |
| Caspase-3 | Cell Signaling # 9662S | 1:1000 |
| GAPDH | EMD Milipore # CB1001 | 1:4000 |
| Cytochrome-C | Cell Signaling #11940S | 1:1000 |
| ATPF1 | Abcam #ab117991 | 1:1000 |
| ATP5A | Abcam, #ab14748 | 1:1000 |
| FLAG tag | Sigma #F7425-.2MG | 1:1000 |

### Cell death assessment (LDH and PI assays)

Neurons were plated at 7.5 x10^4^ cells/well of a tissue culture-treated black 96 well plate coated with laminin and poly-L-lysine with optically clear flat wells 7d prior to experimentation. Propidium iodide or SYTOX green were diluted in complete NBM to a final concentration of 1.5uM or 1uM respectively, and added to neurons during treatment with toxin incubation as described above. Positive control wells were lysed by the addition of 5 ul of 10x lysis buffer (10% v/v Triton X-100) 45 min prior to the desired endpoint for analysis. Propidium iodide uptake was analyzed using the IncuCyte S3 Live cell analysis imaging system and associated software (Sartorius V2018B). To generate time course curves, images were acquired with the IncuCyte ZOOM Plan fluor 20X/0.45 objective (Sartorius cat#4465) every 30 min following PI staining and toxin treatment. PI positive neurons were scored as cells with a threshold signal greater than 2 red calibrated units (RCU) above background, using a Top-hat background subtraction method. Percent maximal uptake was calculated by normalizing experimental wells by the average PI uptake of the triton (positive control) treated wells. For LDH release assays, 50 uL of supernatants from the PI uptake experiments described above were transferred to a clear flat-bottom 96-well plate and assessed using the CytoTox LDH release assay from Promega (cat# G1780) in accordance with the manufacturer’s protocol. Absorbance values (490nm) were obtained on a Bio-Tek Synergy HTX plate reader and analyzed using Gen5 software.

### TMRM assays

Image-iT TMRM (Thermo Scientific cat# I34361) reagent was used at a concentration of 100 nM (1:1000 dilution in NBM). Primary neurons were incubated with TMRM solution for 1h at 37C, and then washed with fresh NBM. Cells were treated with toxins or lentiviruses as described above and imaged every 30 min using the IncuCyte S3 Live Cell Imaging system. Image acquisition and data analysis were performed using the IncuCyte S3 Software (Sartorius V2018B) TMRM positive neurons were scored as cells with a threshold signal greater than 2.5 red calibrated units (RCU) above background, using a Top-hat background subtraction method. Total TMRM intensity was calculated for each well and normalized to DMSO control or vehicle conditions.

### Beta-III-Tubulin (Tuj1) immunocytochemistry

Primary cortical neurons grown on laminin and poly-L-lysine-coated 96-well glass bottom plates were treated with mitochondrial toxins or TDP-43/PR-50 encoding lentiviruses, and stained using standard immunocytochemistry. Briefly, cells were fixed with 4% paraformaldehyde and 15% sucrose in PBS for 1h at RT, rinsed 3 times with PBS, and permeabilized using 0.3% Triton-X-100 in PBS for 1h at RT. Following permeabilization, cells were blocked with 1% BSA in PBS for 1h at RT. Fluorescently conjugated 647-Tuj1 antibody (biolegend cat# 801210) was diluted 1:300 in 1% BSA + 0.3% Triton-X-100 in PBS. Cells were then incubated overnight in Tuj1antibody solution at 4C, and then rinsed 3x with PBS. Cells were then mounted with Prolong Diamond antifade mounting media (Cell Signaling cat# 8961S) and imaged using a Leica Thunder microscope with Andor Zyla sCMOS camera via a 20× Plan Apo objective.

### RNA isolation and qPCR

For analysis of mouse brain tissue, animals were anesthetized with Avertin solution (500 mg/kg, Millipore Sigma) and perfused with 20 mL of cold PBS. Mouse brains were dissected on ice in cold PBS. RNA was then isolated using the RNeasy mini kit (Qiagen). Per the supplier, RNA was isolated by a modified guanidium thiocyanate method and integrity and purity was confirmed using an Agilent 2100 Bioanalyzer. Reverse transcription was performed using the iScript cDNA Synthesis Kit (Bio-Rad). Quantitative real-time PCR (RT-qPCR) was performed using the Power SYBR Green PCR Master Mix (ThermoFisher Scientific) on a StepOnePlus RT PCR system (Applied Biosystems) or a LightCycler 96 (Roche). Expression relative to *Gapdh* was calculated using the comparative C_T_ method.

Primer sequences used for qPCR;

mGSDME F: TGCAACTTCTAAGTCTGGTGACC

mGSDME R: CTCCACAACCACTGGACTGAG *mGapdh*

F: GGGTGTGAACCACGAGAAATATG *mGapdh*

R: TGTGAGGGAGATGCTCAGTGTTG

### Microfluidic chamber assays

For fluidic isolation of local axon segments, neuronal cultures were established in microfluidic devices (RD450 or XC450 from Xona Microfluidics). The silicone microfluidic devices (RD450) were attached onto a glass coverslip (pre-coated with poly-L-Lysine and Laminin) before use.

The microfluidic devices were loaded with freshly dissociated neurons at a density of 500,000 cells per device. Following the initial loading of the neurons into one chamber of each device, they were incubated at 37C for 30 mins to allow the cells to adhere. Afterwards, both chambers of the microfluidic devices were filled with fresh media.The neurons were grown for 9 days in the microfluidic chambers prior to imaging, to allow for sufficient number of axons to grow through the 450µm chamber. During this establishment phase of the neuronal cultures, the growth media in the chamber housing the cell bodies was exchanged with 50% fresh media every 3 days. Neurons grown in microfluidic devices were transfected with lipofectamine 2000 in the same way as described for other cultures. The transfection was carried out in the chamber containing the cell bodies. Transfected neurons were imaged two days following transfection.

### Mouse brain and spinal cord immunohistochemistry

Adult mice were perfused with 20mL of cold PBS and 20mL of 4% PFA. Brains or spinal cords were dissected, fixed overnight in 4% PFA at 4C, paraffin embedded and processed for IHC as described^90^. Briefly paraffin embedded sections were dehydrated in successive washes with xylene and ethanol. Sections were then washed in water and boiled in 1X EDTA unmasking solution (Cell Signaling cat#14746). Tissue sections were then incubated in a 3% H_2_0_2_ solution for 10 min, washed in TBST-Tween20 and blocked for 1h at room temperature with TBST/5% Normal Goat Serum (Cell Signaling Cat#5425). Following blocking, tissue sections were incubated in diluted primary antibody solutions (see below) overnight at 4C. The next day, sections were washed with 1X TBST and incubated for 30min at room temperature in SignalStain^®^ Boost IHC detection Reagent (HRP rabbit, #8114 or HRP mouse, #8125) specific to the species of the primary antibody. Slides were washed with TBST prior to Tyramide Signal Amplification (TSA). Fluorscein conjugated TSA reagent (Akoya Biosciences, NEL745001KT) or Cy3 reagent (Akoya Biosciences, NEL744001KT) were diluted as per manufacturer’s instructions. Slides were incubated with TSA reagent for 10min at room temperature (protected from light). Slides were then washed three times with 1X TBST, counterstained with DAPI and mounted in prolong gold antifade mounting medium (Cell Signaling cat# 8961S) and imaged using a Leica Thunder microscope with Andor Zyla sCMOS camera via a 20× Plan Apo objective. For serial staining of sections (dual IHC) a stripping step was performed by boiling slides for 10 min in a 10mM sodium citrate buffer (Cell Signaling cat# 14746). Following stripping, slides were the incubated in primary antibody solution, washed, incubated with SignalStain Boost IHC secondary detection reagent and subject to TSA amplification as described above.

### Primary antibodies and stains used in mouse brain and spinal cord

Tuj1 (Thermo Scientific, cat # MA1-118X)

GFAP (Cell Signaling, cat# 3670S)

Iba1 (Cell Signaling, cat# 17198S)

CD68 (Cell Signaling, cat# #97778S)

Nissl Neurotrace Green (Thermo Scientific, N21480)

### Human brain immunohistochemistry

Postmortem temporal cortical tissues from FTD/ALS patients with C9orf72 repeat expansions and TDP-43 pathology, patients with Lewy Body Dementia (LBD), and healthy controls were obtained from the UPenn Center for Neurodegenerative Disease Research Biobank. Information on human patients is provided in table S1. Written informed consent was obtained from legal next of kin. Human tissues were processed and stained as previously described ^91^. Briefly, slides were deparaffinized in xylene, and dehydrated in successive EtOH washes. After a brief wash in dH20, slides were then incubated for 30 min in a 70% MeOH and 30% H2O2 solution, and then washed in tap water. Microwave antigen retrieval was performed for 20 min in a citrate buffer (Vector Labs #H-3300). Slides were then cooled to room temperature, rinsed in TBS (0.1 M Tris Buffer) and subject to blocking solution (TBS/2%FBS/3%BSA) for 5 min at room temperature. Following blocking, anti-Human GSDME antibody (abcam 230482, rabbit-anti-human) was diluted 1:100 in blocking solution. Sections were incubated in GSDME primary antibody overnight at 4C in a humidified chamber. The next day, slides were washed with TBS, blocked for an additional 5 minutes and exposed to vector biotinylated anti-Rb IgG secondary antibody (Vector #BA-1000) at a dilution of 1:1000 for 1h at room temperature. Slides were then washed in TBS and Vector ABC solution (Vector # PK-4001) was applied to each slide for 1h at RT. Slides were then exposed to Vector IMPAACT DAB solution at room temperature. Following development, samples were washed in dH20, counterstained using hematoxylin (Cell signaling #14166), rinsed in tap water, dehydrated in successive EtOH and xylene rinses and mounted for imaging.

### Transmission Electron microcopy

Primary mouse neurons were cultured for 7d (as described above). WT and GSDME KO cells were then treated with 5uM raptinal for 1h, prior to processing for electron microscopy. Cells were fixed for at least 2 hours at RT in a 2.5% Glutaraldehyde 1.25% Paraformaldehyde and 0.03% picric acid in 0.1 M sodium cacodylate buffer (pH 7.4) solution. Following fixation, neurons were washed in 0.1M cacodylate buffer and postfixed with 1% Osmiumtetroxide (OsO4)/1.5% Potassium ferrocyanide (KFeCN6) for 1 hour, washed twice with water, once with Maleate buffer (MB), and then incubated in 1% uranyl acetate in MB for 1h followed by 2 water washes and subsequent alcohol dehydration in the following gradations: 10min each; 50%, 70%, 90%, 2×10min 100%. After dehydration propyleneoxide was added to the dish and the cells were lifted off using a transfer pipet, pelleted and infiltrated ON in a 1:1 mixture of propyleneoxide and TAAB Epon (TAAB Laboratories Equipment Ltd, https://taab.co.uk). The following day the samples were embedded in TAAB Epon and polymerized at 60 degrees C for 48 hrs. Ultrathin sections (about 60nm) were cut on a Reichert Ultracut-S microtome, picked up on to copper grids stained with lead citrate and examined in a JEOL 1200EX Transmission electron microscope or a TecnaiG² Spirit BioTWIN and images were recorded with an AMT 2k CCD camera. For analysis, the average mitochondrial length and number of mitochondria per cell was measured using ImageJ. These parameters were compared/normalized to values from WT and KO neurons treated with DMSO.

### Data mining of RNA-seq databases

Single cell transcriptomic data from mouse cortical and hippocampal neurons was obtained from the Allen Brain atlas: https://celltypes.brain-map.org/rnaseq/mouse_ctx-hpf_10x?selectedVisualization=Heatmap&colorByFeature=Cell+Type&colorByFeatureValue=Gad1. Human neuronal single cell data from cortex and hippocampus was obtained and mined from the Allen Brain atlas: https://portal.brain-map.org/atlases-and-data/rnaseq/human-m1-10x. RNA-seq data of FACS-isolated microglia was obtained from the Tabula Muris database co-hosted by the Chan-Zuckerberg Biohub and Broad Institute: https://tabula-muris.ds.czbiohub.org/

### Data mining of postmortem sALS motor neurons

FASTQ files were obtained through the NCBI GEO database (GSE76220) ^56^ and aligned to GRCh38 using STAR (2.7.3a). Differential expression was calculated using DESeq2 (1.36.0) in R (4.0.3). We used a model of design=∼Patient_Sex + Disease to normalize for effects from patient sex.

### Culture, differentiation, transduction, and stressing of human non-disease and ALS patient-specific induced pluripotent stem cell derived motor neurons

Human induced pluripotent stem cells (iPSCs) were generated as previously described (PMID: 25303535, 24507191). IPSCs were cultured on Matrigel-coated (Cat. No. BD354277, VWR) tissue culture plates in StemFlex medium (Cat. No. A3349401, Life Technologies) supplemented with Pen-Strep (Cat. No. 15140163, Life Technologies), maintained at 37°C and 5% CO_2_. Cells were differentiated into motor neurons via chemical-driven, embryoid body (EB)-based protocol as previously described (PMID: 25383599). Briefly, iPSCs were dissociated and grown in suspension culture in StemFlex medium. After 24 hours, cells were grown in N2B27 media consisting of DMEM/F12 (Cat. No. 12634028, Life Technologies) and Neurobasal media (Cat. No. 21103049, Life Technologies) (1:1), N2 supplement (1%) (Cat. No. 17502048, Life Technologies), B27 supplement (2%) (Cat. No. 17504044, Life Technologies), Glutamax (1%) (Cat. No. 35050079, Life Technologies), β-mercaptoethanol (0.1%) (Cat. No. 21985023, Life Technologies), ascorbic acid (20 µM) (Cat. No. A4403, Sigma Aldrich), and Pen-Strep (1%), with additional supplements based on day of differentiation. On days 0 and 1, cells were fed with N2B27 media supplemented with SB 431542 (10 µM) (Cat. No. 1614, R&D Systems), LDN 193189 (100 nM) (Cat. No. 04-0074-02, ReproCELL), and CHIR 99021 (3 µM) (Cat. No. 04-0004-10, ReproCELL). On day 2, cells were fed with N2B27 media supplemented with SB 431542 (10 µM), LDN 193189 (100 nM), CHIR 99021 (3 µM), retinoic acid (1 µM) (Cat. No. R2625, Sigma Aldrich), and smoothened agonist (SAG) (1 µM) (DNSK International). On day 4, media was replaced with day 2 media. On day 5, media was replaced with N2B27 media with retinoic acid (1 µM) and SAG (1 µM). On day 7, cells were fed with N2B27 media with RA (1 µM), SAG (1 µM), and BDNF (20 ng/mL) (Cat. No. 248-BD, R&D Systems). On day 9, media was replaced with N2B27 supplemented with RA (1 uM), SAG (1 uM), BDNF (20 mg/mL) and gamma secretase inhibitor (DAPT, 10 µM) (Cat. No. 2634, R&D Systems). On day 11, cells were fed with N2B27 with RA (1 µM), SAG (1 µM), DAPT (10 µM), BDNF (20 ng/mL), and GDNF (20 ng/mL) (Cat. No. 212-GD, R&D Systems). On day 13, media was replaced with day 11 media. On day 15, EBs formed were dissociated with 0.25% Trypsin-EDTA (Cat. No. 25200114, Life Technologies, counted, and centrifuged at 400 x g for 5 minutes. Pelleted cells were resuspended in complete MN media consisting of Neurobasal media, N2 supplement (1%), B27 supplement (2%), Pen-Strep (1%), Glutamax (1%), nonessential amino acids (1%) (Cat. No. 11-140-050, Gibco), βME (0.1%), ascorbic acid (20 µM), BDNF (20 ng/mL), GDNF (20 ng/mL), CNTF (20 ng/mL) (Cat. No. 130-096-336, Miltenyi), and UFDU (10 µM) (Cat. Nos. U3750/ F0503, Sigma Aldrich). Plated cells were cultured for an additional 7 days prior to experimentation, with half media changes every 3-4 days.For GSMDE knockdown experiments, MNs were plated at 100,000 live cells/well in a 96-well plates pre-coated with poly-L-ornithine (Cat. No. P3655, Sigma Aldrich), poly-D-lysine (Cat. No. A3890401, Life Technologies), fibronectin (Cat. No. 47743-654, VWR), and laminin (Cat. No. 23017015, Life Technologies). Cells were infected with an MOI of 6 and cultured for 7 days, with half media changes every 3-4 days. After 7 days, transduced cells were exposed to thapsigargin (0.5 µM) (Cat. No. T9033, Sigma Aldrich), tunicamycin (5 µM) (Cat. No. T7765, Sigma Aldrich), or MG132 (1 µM) (Cat. No. M8699, Sigma Aldrich) for 48 hours and subsequently fixed with 4% PFA.

### Data analysis and code availability

Enrichment analysis of GSDME on mitochondria, quantification of GSDME puncta and analysis of microtubule depolymerization was done using custom Fiji (ImageJ) macros (available at: https://github.com/IsaacChiuLab/FIJIanalysisCodes_GSDME).

#### Colocalization analysis of GSDME and mitochondria along neurites

For analyzing the enrichment of GSDME on mitochondria along neurites, the neurites were tracked manually. The mitochondrial intensity and the GSDME intensity were then measured along the neurites. The mitochondrial intensity was then converted to 1-D masks (by thresholding according to local contrast) while the GSDME intensity was normalized to the average intensity along the neurite. The enrichment of GSDME was then calculated as the ratio of mean GSDME signal within the mitochondrial mask to the mean GSDME signal outside the mitochondrial masks.

#### Colocalization analysis of GSDME and mitochondria in cell bodies and SHSY5Y cells

To analyze the enrichment of GSDME on mitochondria in cell bodies and SHSY5Y cells, the cell outline was drawn manually. The mitochondrial intensity within the cell outline was then used to create masks (by thresholding according to local contrast) and the GSDME intensity was normalized to the average intensity of the whole cell. As done in the case of neurites, enrichment of GSDME on mitochondria was reported as the ratio of mean GSDME signal within the mitochondrial mask to the mean GSDME signal outside the mitochondrial masks

#### Quantification of GSDME puncta

To quantify the density of puncta along neurites, the neurites were manually traced. The intensity of GSDME was then measured along the neurites and normalized to the average intensity along the whole neurite. To detect GSDME puncta, the GSDME signal was then converted to a mask using a local thresholding window of 20px. The density of GSDME puncta in the cell bodies and SHSY5Y cells was quantified using a similar approach. In this case, the cells were traced manually, following which the GSDME puncta were detected by local thresholding.

#### Analysis of Beta-III-Tubulin (Tuj1) immunocytochemistry

The microtubule (MT) depolymerization index was measured using a custom macro adapted from ^39^. Briefly, the polymerized microtubules were distinguished from depolymerized ones using brightness and circularity. Polymerized microtubules are less bright and less circular than the aggregated puncta formed by depolymerized microtubules. In this macro, a low threshold is set at first, to detect and mask all TUJ1 positive objects (including polymerized and depolymerized MTs). In the second step, a high threshold is set, which can distinguish only the bright spots formed by depolymerized MTs. Once detected and masked, the pixels marking depolymerized MTs are removed from the set of pixels (mask) marking all microtubules, as detected in the first step, to get the mask of pixels containing polymerized microtubules only. In the final step, the masks of polymerized and depolymerized microtubules are then refined using a circularity threshold. Any tubular objects detected in the mask of depolymerized microtubules or any solid round objects detected in the mask of polymerized microtubules are removed. The MT depolymerization index is then measured as the following ratio:

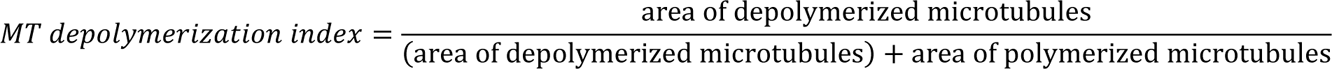

All thresholds used for MT depolymerization analysis were determined prior to analyzing full datasets, using negative and positive controls.

#### Quantification of mitochondrial motility in neurites

To quantify mitochondrial movement in neurites, images of fluorescently marked mitochondria were captured every 6s. Kymographs were generated from 3 to 5-min time-lapse movies and analyzed with Kymolyzer ^87^, a custom-written ImageJ macro. Each data point in the percent motility quantifications represent the average percent time spent in motion by all mitochondria in a neurite segment.

## Supplemental Figure Legends

**Figure S1: GSDME is expressed in mouse and human neurons (related to Figure 1 & 2)**

(A) *Gsdme* RT-qPCR of several mouse brain regions from WT (n=3) and *Gsdme* knockout (n=2) mice

(B) Representative IHC images of cortical mouse brain sections from WT and GSDME knockout animals stained with anti-GSDME and counterstained with DAPI.

(C) FACS-isolated brain myeloid cell transcriptomic data from the publicly available Tabula Muris database was mined for *Gsdme* and *Gsdmd* expression. TSNE plots showing cell types (left most plot) and expression of gasdermin genes (right) were generated using the CZI biohub tool (Available from: https://tabula-muris.ds.czbiohub.org/)

(D) Publicly available human cortical single-cell RNA-seq (10X Genomics) data was mined for gasdermin family expression levels. TSNE plots showing cell types (left most plot) and gasdermin genes were generated using the Allen Brain Atlas Transcriptomics Explorer tool (Allen Institute for Brain Science. Available from: <u>human.brain-map.org</u>)

(E) SH-SY5Y cells were incubated in propidium iodide (PI) containing media and treated with 5uM raptinal, 20uM rotenone or 30uM of 6-OHDA. Images were taken every 3h and quantified for PI uptake.

(F) primary neurons were incubated in propidium iodide (PI) containing media and treated with 5uM raptinal, 20uM rotenone and subsequently imaged to quantify PI uptake.

(G) SH-SY5Y were treated with 5uM raptinal, 20uM rotenone or 30uM of 6-OHDA and assessed for LDH release at 24h post-treatment.

(H) Primary neurons were incubated in media containing multiple doses of raptinal or rotenone and assessed for LDH release at 24h post-treatment.

**Figure S2: GSDME is activated by mitochondrial and ER toxins and mediates cell death in human SH-SY5Y cells (related to Figure 2).**

(A) Immunoblots of WT and GSDME KO SH-SY5Y treated with DMSO, raptinal, rotenone or 6-OHDA. Cells treated with raptinal were harvested for lysates at 2h post-treatment, while cells treated with rotenone or 6-OHDA were collected at 12h. All concentrations are in micromolar units.

(B) Representative 20X Images of WT (left) or GSDME KO (right) SH-SY5Y cells at 2h following treatment with 5uM raptinal. Cells were incubated in *Sytox* Green-containing media, and green cells indicate dye uptake. Scale bar: 200 um

(C-G) WT SH-SY5Y and GSDME KO cells treated with (C) 5uM raptinal (D) 20uM rotenone (E) 30uM 6-OHDA (F) 1uM antimycin or (G) 1mM 3-nitropropionic acid were co-incubated with propidium iodide, imaged every 3h and quantified for PI uptake.

(H) WT and GSDME KO SH-SY5Y were treated with 5uM raptinal (1h), 20uM rotenone (4h) or 30uM 6-OHDA (12h) were assessed for LDH release at 24h post-treatment.

(I) Representative 20X images of GSDME wild-type or GSDME knockout SH-SY5Y cells stained with propidium iodide (PI) and treated with DMSO or 0.4uM thapsiagargin (Tg). Scale bar: 200um (J-K) SH-SY5Y were incubated in propidium iodide containing media and treated with (G) 0.4uM thapsiagargin (Tg) or (H) 4uM tunicamycin. Images were taken every 4h in an IncuCyte ZOOM machine and propidium iodide uptake was quantified.

**Figure S3: Caspase-3 but not RIPK1 inhibition protects neurons from mitochondrial toxins (related to Figure 2)**

(A-B) Immunoblots of (A) primary mouse neurons or (B) SH-SY5Y treated with DMSO or raptinal (5-10uM) in the presence or absence of 20uM zVAD-FMK.

(C-D) Representative 20X images of primary neurons stained with PI and treated with either 5uM raptinal or 5uM raptinal + 20uM zVAD-FMK (C). Images were taken every 3h by an IncuCyte ZOOM and (D) PI uptake was quantified. Scale bar: 200um

(E-F) Representative 20X images of SH-SY5Y stained with PI and treated with either 5uM raptinal or 5uM raptinal + 20uM zVAD-FMK (E). Images were taken every 3h in an IncuCyte ZOOM and

(F) PI uptake was quantified. Scale bar: 200um

(G-H) Primary mouse neurons treated with (G) a5uM raptinal or (H) 30uM rotenone were co-incubated in media containing propidium iodide and either DMSO, nec1 or GSK ‘872 (RIPK1 inhibitors). Images were taken every 3h and quantified for PI uptake.

**Figure S4: GFP-GSDME cleavage by mitochondrial toxins and downstream caspase-3 causes enhanced cell death and puncta formation (related to Figures 2 & 3)**

(A) Diagram showing the full-length human GSDME (yellow), and caspase-3 cleavage site (dotted line). An eGFP (green) sequence was placed on the N-terminal side of hGSDME.

(B-C) HEK 293T cells were transfected with GFP (control) or GFP-fl-GSDME and treated with either DMSO or raptinal (5uM). Immunoblots for (B) GSDME, (C) GFP or CASP3 show cleavage and activation of CASP3 and the GFP-GSDME construct.

(D-F) Cells were transfected with GFP-GSDME (top panels) or control vector (bottom panels) and co-incubated with PI and either raptinal or rotenone. Cells were imaged every 1h and (E-F) quantified for PI uptake and GFP expression.

(G) Primary WT neurons transfected with GFP-GSDME were incubated with 5uM raptinal and immunoblots were performed with cells treated for different times to assess the temporal dynamics of caspase-3 and GSDME activation.

**Figure S5: Mitochondrial toxins induce GFP-tagged GSDME to form puncta in a caspase-dependent manner (related to Figures 3).**

(A-B) SH-SY5Y cells treated with raptinal + DMSO, or raptinal + 20uM zVAD, and imaged over 90 minutes. White arrows represent intracellular GFP-GSDME puncta, while arrowheads denote plasma membrane enrichment (A) Images were captured every 10 min and the frequency of cells showing intracellular puncta, as well as positive PI staining were (B) quantified in the (top) absence or (bottom) presence of 20uM zVAD. Scale bar: 10um

(C) Primary mouse neurons were co-transfected with mTagRFP-Membrane-1 (neuromodulin: aa 1-20) and GFP-GSDME and treated with 5uM raptinal. Following toxin treatment, images were taken every 30 min (for 10h total) in order to look for colocalization of these two markers. Scale bar: 20um

(D) Primary neurons transfected with GFP-GSDME were treated with 5uM raptinal or raptinal+zVAD-FMK and imaged 45 minutes after drug treatment. Top (large) panels show representative single neurons prior to any drug treatment. Scale bars: 50 um. The bottom panels show magnified axonal segments (white boxes) before and after 45 min of drug treatment. Scale bars: 20 um

(E) Quantification of GFP-GSDME puncta treated with either 5uM raptinal or 5uM raptinal + 20uM zVAD-FMK. Each dot represents a single neuron, and 8-10 neurons from three independent experiments were used for quantification. Data was analyzed using paired Student’s t-tests comparing before and after drug treatment. P-values were adjusted for multiple comparisons using the Tukey method.

**Figure S6: Toxin treatment causes GSDME to colocalize with mitochondria (related to Figure 3)**

(A-B) Representative cell bodies from (A) mouse neurons transfected with mitochondrial marker mKate-OMP25 (red) and GFP-GSDME (green) before and after 3h treatment with 5uM raptinal are shown. (B)The enrichment of the green signal on mitochondria over cytosol was quantified and compared before and after raptinal treatment (N=12 cell bodies representing neurons from 3 replicate experiments).

(C-E) Mouse neurons transfected with mitochondrial marker mKate-OMP25 or GFP-GSDME were imaged before and after treatment with 20uM rotenone. (C) Images were captured 4h post-rotenone treatment. (D) The location of red (mitochondria) and green (GFP-GSDME) peaks were identified along the neurites. (E) The intensity of the red signal overlapping with green GSDME peaks was quantified and compared before and after raptinal treatment.

(F) SH-SY5Y transfected with mitochondrial marker mKate-OMP25 or GFP-GSDME were imaged before and after treatment with 5uM raptinal. Images were captured 2.5h post-raptinal treatment. (G-H) Immunoblot of enriched cytosolic and mitochondrial (ATP5F1 positive) fractions from SH-SY5Y treated with either DMSO or 10uM raptinal for 1h. (G) The top two panels represent full length and cleaved GSDME fragments. (H) Quantification of the intensity of cleaved over full length GSDME bands in immunoblots from cytosolic or mitochondrial SH-SY5Y fractions (n=3). (I-J) Immunoblot of enriched cytosolic and mitochondrial (ATP5F1 positive) fractions from primary neurons treated with either DMSO or 10uM raptinal for 1h. The top two panels (I) represent full length and cleaved GSDME fragments. (J) Quantification of the intensity of cleaved over full length GSDME bands in immunoblots from cytosolic or mitochondrial primary neuron fractions (n=3)

(K-L) Immunoblots of WT or GSDME KO SH-SY5Y treated with DMSO or 5uM raptinal for 1h. (K) Mitochondrial (ATP5F1) and cytosolic fractions were blotted for GSDME, cl-casp3 and cytochrome-c (cyt-c). The ratio of mitochondrial to cytosolic cytochrome-c signal by immunoblot was (L) quantified for WT and KO cells treated with 5uM raptinal (n=4).

**Figure S7: GSDME recruitment to mitochondria is cardiolipin dependent (related to Figure 3)**

(A) Primary mouse cortical neurons were transduced with lentiviruses encoding 3 different shRNAs (#1-3) targeting mouse cardiolipin synthase 1 (CLS1) and scrambled control. Immunoblotting against CLS1 revealed efficient KD with shRNA#2

(B-C) Representative images of GSDME KO mouse neurons transduced with lentivirus encoding CLS1-shRNA#2 or scrambled control and transfected with both GFP-GSDME and OMP25-mKate. 3d post-transduction, cells were treated with 5uM of raptinal and imaged at 0, 3h and 4h post treatment. (C) Enrichment of GFP-GSDME puncta colocalized with mitochondria were quantified at baseline, 3h and 4h post-raptinal treatment.

**Figure S8: Toxins cause GSDME activation and mitochondrial colocalization in human iNeurons (related to Figure 3).**

(A-B) Immunoblots of human iPSC-derived cortical neuron (iNeuron) cultures. (A) Immunoblot of human iNeurons treated with 5 or 10uM raptinal for 1h. (B) Immunoblot of human iNeurons treated with DMSO, 20uM zVAD, 5uM raptinal or a combination of raptinal and zVAD for 1h.

(C-E) Human iPSC-derived cortical neurons (iNeurons) transfected with mitochondrial marker mKate-OMP25 and GFP-GSDME were imaged before and after a 2.5h treatment with 5uM raptinal. A representative neuron (C, top panel), with magnified images of a neurite (indicated by dotted white box) before and after raptinal treatment are shown (C, bottom panel). (D) Locations of mitochondria and intensity of GFP-GSDME were measured as line scans along neurites. Line scans (as shown in panel D) were analyzed to calculate (E) GFP-GSDME enrichment on mitochondria before and after raptinal treatment. N >100 axon segments representing iNeurons originating from 3 independent differentiations.

**Figure S9: GSDME deficiency protects from TMRM loss in primary neurons and SH-SY5Y (related to Figure 4)**

(A-D) Wild-type and GSDME knockout primary cortical neurons were stained with TMRM and incubated with (A-B) 0.5 and 1uM raptinal or (C-D) 10uM and 20uM Antimycin-A. Representative images of TMRM uptake are shown (A and C). TMRM intensity was quantified at 30 min and 1h post-toxin treatment, respectively (B and D).

(E-G) Wild-type and GSDME knockout SH-SY5Y were stained with TMRM and incubated with (E-F) 0.5uM or 3uM raptinal or (G) 10uM rotenone. Images captured at 4 and 10h post-treatment

(E) were quantified for TMRM intensity (F-G).

All TMRM intensities are represented as intensities to respective DMSO controls.

**Figure S10: GSDME deficiency protects from caspase-3 activation and neurite loss (related to Figures 4 & 5).**

(A) Representative images at several timepoints of wild-type and GSDME knockout SH-SY5Y cells treated with 2uM raptinal and incubated in media containing Incucyte Caspase-3/7 dye.

(B-C) WT and KO SH-SY5Y incubated in Caspase-3/7 dye were treated with (B) 5uM raptinal and imaged every 3h post-treatment. (C) Area under the curve (AUC) measurements from 5uM raptinal (AUC_6h_), 20uM rotenone (AUC_24h_) and 30uM 6-OHDA (AUC_12h_) treatment were generated and compared between WT and KO cells.

(D-E) WT and KO primary neurons incubated in Caspase-3/7 dye were treated with (D) 5uM raptinal and imaged every 2-3h post-treatment. (E) Area under the curve (AUC_24h_) measurements from 5uM raptinal, 20uM rotenone and 20uM antimycin-A treatment were generated and compared between WT and KO cells.

(F) Representative images of wild-type neurons treated with DMSO, 3uM raptinal and 20uM antimycin-A and stained for Tuj1 at 8h post-toxin treatment.

(G) Microtubule depolymerization index was calculated for WT neurons treated with several doses of raptinal and antimycin-A. Violin plots display the combined median and interquartile ranges for depolymerization index.

(H) WT and GSDME KO mouse cortical neurons were treated with 4uM raptinal, 20uM rotenone or 20uM Antimycin-A and were assessed for LDH release at 8h post-treatment.

(I) Microtubule depolymerization index was calculated for WT neurons co-incubated with either DMSO or 20uM of zVAD-FMK and were then treated with several doses of raptinal, antimycin-A or rotenone (all concentrations are mentioned in micromolar units). Violin plots display the combined median and interquartile ranges for depolymerization indices taken from two independent experiments.

(J) Representative image of a microfluidic chamber plated with wild-type mouse cortical neurons stained with TMRM. The cell body, microgrooves and axonal compartments are labeled. The top panels represent the chamber prior to addition of 5uM raptinal, while the bottom panels show 1h post-toxin treatment.

(K) Quantification of TMRM intensity relative to baseline (time = 0) from the cell body and axonal chambers of plated WT neurons treated with 5 or 10uM raptinal.

For all datasets two-way ANOVA (row factor = toxin, column factor = genotype) was performed, followed by multiple comparisons for each group (p-values adjusted by the Tukey method are mentioned over respective comparisons). Data represents an average of at least 2 independent experiments.

**Figure S11: Expression of N-GSDME in neurons is sufficient to drive mitochondrial localization, mitochondrial damage and neurite loss (related to Figure 5)**

(A) Representative widefield images of mouse neurons transfected with mKate2-OMP25 and either full-length GFP-GSDME (top row), N-terminal GSDME (middle row) or untransfected control (bottom row). Cells were imaged 16h post-transfection.

(B) Structured illumination microscopy (SIM) images of mitochondria (OMP25-mKate) in axons of mouse neurons transfected with GFP-N-GSDME to assess colocalization.

(C) Representative widefield images of SH-SY5Y transfected with mKate2-OMP25 and either full-length GFP-GSDME (bottom image) or GFP-N-GSDME (top image).

(D) Structured illumination microscopy (SIM) images of SH-SY5Y mitochondria (OMP25-mKate) transfected with GFP-N-GSDME to assess colocalization.

(E-F) Primary mouse neurons (DIV3) were transduced with lentivirus encoding either N-GSDME or GFP-control. (E) 4d post-transduction (DIV7) neurons were stained with TMRM. (F) Images from 6 wells per condition across 2 independent experiments were quantified by measuring TMRM intensity along the length of neurites.

(G-H) Primary mouse neurons (DIV3) were transduced with lentivirus encoding either N-GSDME or GFP-control. (G) 4d post-transduction, neurons were fixed and stained for Tuj1+ signal. (H) Images from 6 wells per condition across 2 independent experiments were used to calculate microtubule depolymerization index for each condition.

For datasets F, H one-way ANOVA was performed, followed by multiple comparisons for each group (adj p-values, Tukey method). For F,H data represents an average of 2 independent experiments (3 technical replicates/experiment) ± SEM.

**Figure S12: GSDME is activated in iNeurons and mediates neurite loss caused by FTD/ALS proteins (related to Figure 6)**

(A) GSDME immunoblot quantification (n=5-6) of primary cortical neurons transduced with lentivirus encoding GFP, PR-50, TD-43 or un-transduced control. Four days following transduction with lentiviruses, samples were lysed and probed with anti-GSDME and GAPDH (loading control). The ratio of N-terminal GSDME to full-length was calculated and plotted relative to untreated control.

(B) Representative images of WT and GSDME KO mouse cortical neurons stained with TMRM (red) and transduced with lentiviruses encoding PR-50 or TDP-43 (4d post-transduction). Scale Bar: 200um

(C-E) Human iNeurons were transfected with mKate-OMP-25, GFP-GSDME and TDP-43 or iRFP (control). (C) Cells were imaged 72h post-transfection (D) Quantification of the number of GFP-GSDME puncta per 100um in neurites.

(E) Human iNeurons were transduced with control virus (GFP) or lentivirus encoding FLAG-TDP43 at multiplicity of infection 4 or 8. 5d post-transduction, cells were lysed for immunoblot analysis.

(F-I) Primary WT and GSDME knockout mouse neurons were co-transfected with Syn1-RFP and either Syn1-GFP, (F-G) Syn1-TDP-43 or (H-I) Syn1-PR-50 and imaged for neurite (RFP signal) area. Representative images of WT and KO neurons transfected with Syn1-RFP and either TDP-43 (F) or PR-50 (H) are shown along with the quantifications of neurite area (G and I). Each dot represents an average of 10-12 transfected neurons imaged in an independent well. Every condition was imaged across two independent experiments.

For datasets A, G, I two-way ANOVA (row factor = lentivirus treatment, column factor = genotype) was performed, followed by multiple comparisons for each group (adj p-values, Tukey method). For F-I, data represents an average of 2 independent experiments ± SEM.

**Figure S13: GSDME is activated in the SOD1 G93A mouse model and plays a role in disease progression (related to Figure 7)**

(A-B) GSDME and GAPDH immunoblots of spinal cord lysates from (A) pre-symptomatic stage (P82) and end-stage (P160) SOD1^G93A^ mice. (B) The activation ratio of GSDME is quantified as the levels of GSDME N-terminal compared to total GSDME (full length + cleaved fractions).

(C) Expression of the SOD1^G93A^ transgene, as assessed by RT-qPCR, was quantified for each group and genotype.This was done to confirm that the copies of this ALS-causing transgene were comparable between experimental groups.

(D) Weights of transgenic SOD1^G93A^ GSDME WT or SOD1^G93A^ GSDME KO mice were tracked over time starting at week 7 until week 21.

(E) Grip strength measures of nTg WT and nTg GSDME KO mice (lacking SOD1^G93A^) were recorded at P60 and P150 and normalized by body weight.

(F) Table of summary values for male and female transgenic mice displaying key measures of survival, disease progression and motor function.

**Figure S14: GSDME knockout reduces histological markers of gliosis in the SOD1^G93A^ mouse model of ALS (related to Figure 7)**

(A-B) Representative images of lumbar spinal cord sections from SOD1^G93A^ GSDME WT and SOD1^G93A^ GSDME KO animals at P150 (A) stained with anti-GFAP to mark astrocytes. (B) The intensity of GFAP in the ventral horn of P150 transgenic mice normalized by the ventral horn area was quantified. Each dot represents the average value of 6-8 stained lumbar spinal cord sections taken from a single mouse. Magnified regions (white box) delineate a ventral horn area of a spinal cord section. White scale bars (large images) = 200um.

(C-D) Representative images of lumbar spinal cord sections from SOD1^G93A^ GSDME WT and SOD1^G93A^ GSDME KO animals at P150, stained with anti-Iba1(C). Magnified regions represent the ventral horns of the spinal cord used for (D) quantification of mean Iba1 counts per um^2^. White scale bars (large images) = 200um.

(E-F) Representative images of lumbar spinal cord sections from SOD1^G93A^ GSDME WT and SOD1^G93A^ GSDME KO animals at P150, stained with anti-CD68 (a lysosomal activation marker), to assess disease associated inflammation (E). Magnified regions represent the ventral horns of the spinal cord used for (F) quantification of mean CD69 counts per um^2^. White scale bars (large images) = 200um.

**Figure S15: Endoplasmic reticulum stressors and proteasome inhibition cause neurite loss in susceptible TDP43 G298S iPSC-derived motor neurons (related to Figure 8)**

(A) RNA-seq transcriptomic data from laser capture micro-dissected motor neurons from patients with sporadic ALS and age-matched controls (Krach et al., 2018)

(B) Representative images of control 1016A or *TDP43*^G298S^ iPSC-derived motor neurons treated with DMSO, MG132, tunicamycin or thapsiagargin, and stained for Tuj1 at 48h post-toxin treatment.

(C-E) Microtubule depolymerization index was calculated for 1016A (WT) and *TDP43*^G298S^ motor neurons treated with DMSO (vehicle) or several doses (C) MG132 (D) thapsiagargin or (E) tunicamycin (uM). Violin plots display the combined median and interquartile ranges for depolymerization index.

(F-G) Immunoblots of iPSC-derived cortical neurons 4d post-transdution with scrambled or GSDME-targeting shRNAs(F), which was quantified in (G).

## Supplementary videos

**Supplementary video 1:** Live cell imaging of GFP-GSDME transfected SH-SY5Y cells treated with 5uM of raptinal and either DMSO (top) or 20uM of zVAD-fMK (bottom). Cells were imaged every 10 minutes and GFP (GSDME) and RFP (PI) channels were collected.

**Supplementary video 2-3:** Representative reconstructions of mitochondria from mouse neurons transfected with GFP-GSDME and either (video 2) OMP25-mCherry or (video 3) Cox8-mCherry. These cells were treated with 10uM raptinal were fixed and imaged using structured illumination microscopy (SIM) to assess marker colocalization.

**Supplementary video 4:** Recordings of female (top row) and male (bottom row) of SOD1^G93A^ transgenic mice that are either GSDME KO (left panels) or GSDME WT (right panels). All recordings were taken from mice at P150.

